# Spatially-restricted metabolic delivery of auxin for selective adventitious root induction

**DOI:** 10.64898/2026.08.20.746056

**Authors:** Qianqian Liu, Yinwei Zeng, Sebastien Schotte, Damilola Olatunji, Markéta Luklová, Ren Wang, Hoang Khai Trinh, Geert Goeminne, Karin Ljung, Ondrej Novak, Federica Brunoni, Ohad Roth, Roy Weinstain, Jonas Mortier, Thomas Heugebaert, Inge Verstraeten, Danny Geelen, Steffen Vanneste

**Affiliations:** HortiCell, Department of Plants and Crops, Faculty of Bioscience Engineering, Ghent University, Ghent, Belgium; Department of Plant & Microbial Biology, University of California, Berkeley, CA 5 94720, USA; Department of Genetics, Development and Cell Biology, Iowa State University, Ames, IA 50011; VIB Metabolomics Core, Ghent, Belgium; Department of Plant Biotechnology and Bioinformatics, Ghent University, Ghent, Belgium; Department of Forest Genetics and Plant Physiology, Umeå Plant Science Centre, Swedish University of Agricultural Sciences, 901 83 Umeå, Sweden; Laboratory of Growth Regulators, Faculty of Science, Palacký University Olomouc and Institute of Experimental Botany of the Czech Academy of Sciences, Olomouc, Czech Republic; Department of Biotechnology, University of Verona, Verona, Italy; School of Plant Sciences and Food Security, Faculty of Life Sciences, Tel Aviv University, Tel Aviv, Israel; Research Group SynBioC, Department of Green Chemistry and Technology, Ghent University, Ghent, Belgium; HortiRoot, Department of Plants and Crops, Faculty of Bioscience Engineering, Ghent University, Ghent, Belgium

## Abstract

Auxin impacts on nearly every aspect of plant growth and development. Its exogenous application therefore results in pleiotropic growth responses. Exploiting this activity for plant propagation requires avoiding or minimizing such off-target effects and is generally achieved as a trade-off between toxicity and organogenetic efficacity. We recently identified the compound HYSPARIN (HYS) with potent, and uniquely selective adventitious root inductive activity. Unlike other root-inducing compounds, HYS preferentially activates auxin responses in the shoot via an unknown mechanism. Here, we show that HYS acts as a shoot-specific proauxin. Rather than acting through auxin homeostasis, we found that HYS is hydrolysed i*n planta* independently of ILR1/ILL amidohydrolases to release the synthetic auxin MCPA. Structure–activity relationship analysis confirmed a strong dependence on its MCPA moiety for activating auxin responses, and identified its promoiety as a determinant of shoot-specificity and activity. Selective application of MCPA also potently induces AR is consistent with a model in which HYS metabolism produces a spatially restricted, AR inductive auxin signal. The activation mechanism of HYS thus provides a conceptual framework for tissue-specific metabolic delivery of auxin and may enable the programmable delivery of other xenobiotics in plants.

## Introduction

While auxin has many important functions in stimulating plant growth and development, its exogenous application readily becomes herbicidal due to the activation of uncontrolled, systemic auxin responses (Grossmann, 2010). Despite this limitation, auxin is also commonly used to induce roots in stem cuttings during clonal propagation of many species (Blythe et al., 2007).

In shoot cuttings, endogenous auxin accumulates close to the wound as a consequence of disruption of the auxin flow (Bellini et al., 2014). In easy-to-root species, this auxin signal is sufficiently strong to trigger adventitious rooting. In recalcitrant species or recalcitrant, mature tissues, an additional dose of auxin can help to improve rooting efficiency (Blythe et al., 2007). The synthetic auxin analogues are typically more stable than IAA, and are therefore often used in rooting mixtures (Blythe et al., 2007), but can be phytotoxic (Busi et al., 2018).

Indole-3-butyric acid (IBA) is a well-established adventitious root inducer (Zimmerman and Wilcoxon, 1935). It can release bioactive IAA upon beta-oxidation in the peroxisomes (Damodaran and Strader, 2019), providing a metabolic gate to its activity. The advent of chemical screening in plants has identified several novel AR-inductive molecules. Arinole is a benzoxazole that indirectly induces AR formation in multiple species by stimulating IAA biosynthesis (Depaepe et al., 2024). The tryptophan-conjugated synthetic auxin, 4-chlorophenoxyacetic acid-L-tryptophan-O-me (named 1q) is metabolically hydrolysed to auxin, and helps to overcome AR recalcitrance in woody species, such as apple and argan (Roth et al., 2024). We previously identified a small molecule that potently induces AR in hypocotyls of etiolated seedlings (Zeng et al., 2023). HYS has limited to no effects on root growth and development, a feature that is unique among AR-inducers. It is currently unclear what determines its shoot-specific activity.

Proauxins, like IBA and 1q, are auxin analogs that are inactive until metabolized. This can be either by modifying a functional group (bioprecursors), as exemplified in the beta-oxidation of the carboxylgroup in IBA or by the enzymatic hydrolysis of a linked promoiety that keeps the auxin inactive (carrier-linked proauxins), as exemplified by conjugated auxins, such a 1q. The metabolically gated hydrolysis and resistance to homeostatic clearance makes these proauxins persistent, slow-release pools of auxin (Roth et al., 2024). This activity profile makes proauxins effective tools for stimulating hypocotyl elongation, adventitious rooting or lateral root development (Savaldi-Goldstein et al., 2008; Kerchev et al., 2015; Roth et al., 2024).

Members of INDOLE-3-ACETIC ACID-LEUCINE RESTANT1/ILR1-LIKE (ILR1/ILL) amido hydrolase family, ILR1 and INDOLE-3-ACETIC ACID-ALANINE RESISTANT3 (IAR3) have been identified as enzymes capable of releasing free IAA from amino acid conjugated IAA (Bartel and Fink, 1995; Davies et al., 1999). Five additional *ILL*s were identified in the Arabidopsis genome of which ILL2 has a strong catalytic activity towards IAA conjugates (LeClere et al., 2002). The conjugated amino acid severely impacts on the hydrolysis rates effected by the different ILR1/ILLs, indicating an important degree of substrate specificity. Interestingly, their activity range is not restricted to IAA conjugates, but extends to amino acid conjugates of oxIAA (Hayashi et al., 2021), Phenyl Acetic Acid (PAA) (Hladík et al., 2025), Jasmonic Acid (Widemann et al., 2013; Zhang et al., 2016), and its precursor 12-oxo-phytodienoic acid (OPDA) (Široká et al., 2024). Recently, ILR1/ILLs were also found to be active towards the demethylated derivative (1r) of the Trp-conjugated proauxin 1q (Roth et al., 2024), and Asp- or Glu-conjugated 2,4-D (Chiu et al., 2018). This broad activity range suggests that ILR1/ILL enzymes exhibit substrate promiscuity towards amino acid-conjugates.

We recently identified HYSPARIN, a small molecule that induces AR in hypocotyls by selective activation of auxin signaling in the shoot via an unknown mechanism (Zeng et al., 2023). Here, we explored the determinants of the selective activity of HYSPARIN. We could exclude that HYS AR-induction acts through modulation of endogenous auxin biosynthesis or catabolism. Instead, we show that HYS acts as a proauxin that is metabolized independent of ILR1/ILL amido hydrolase activity to deliver the synthetic auxin 2-methyl-4-chlorophenoxyacetic acid (MCPA) to the shoot. Structure-activity relationship confirmed that HYS auxinic activity resides in its MCPA moiety, while the tissue-specificity largely resided in the anilide-containing promoiety. Jointly, our work identifies HYS as a tissue-specific proauxin that enables the targeted metabolic delivery of auxin to the shoot for AR induction.

## Results

### Endogenous IAA biosynthesis supports but does not mediate HYS-induced AR formation

First, we interrogated the involvement of endogenous auxin homeostasis in HYS’ ability to induce AR. Mutants defective in Trp-biosynthesis (*wei2-1*, and *wei2-1wei7-1*) and at the first step of the subsequent conversion to IAA (*wei8-1*, *wei8-1tar2-1*, *wei8-2tar2-2*), developed fewer AR in response to HYS (Fig. 1A; Supplementary Fig. S1A). Consistently, treatment with the IAA biosynthesis inhibitor L-kynurenine (KYN) (He et al., 2011), which inhibits TAA1/TAR activity significantly reduced HYS-induced AR formation (Fig. 1B). Mutants (*ech2-1ibr10-1*) in IBA to IAA conversion also displayed a reduced HYS response (Figs. 1C; Supplementary Fig. S1B). In contrast, reduced auxin inactivation in *gh3* octuple (*gh3.1-6/9/17*) mutants was associated with increased AR formation (Figs. 1D; Supplementary Fig. S1C). These data reveal that auxin homeostasis impacts on HYS-induced AR formation.

**Figure 1.**
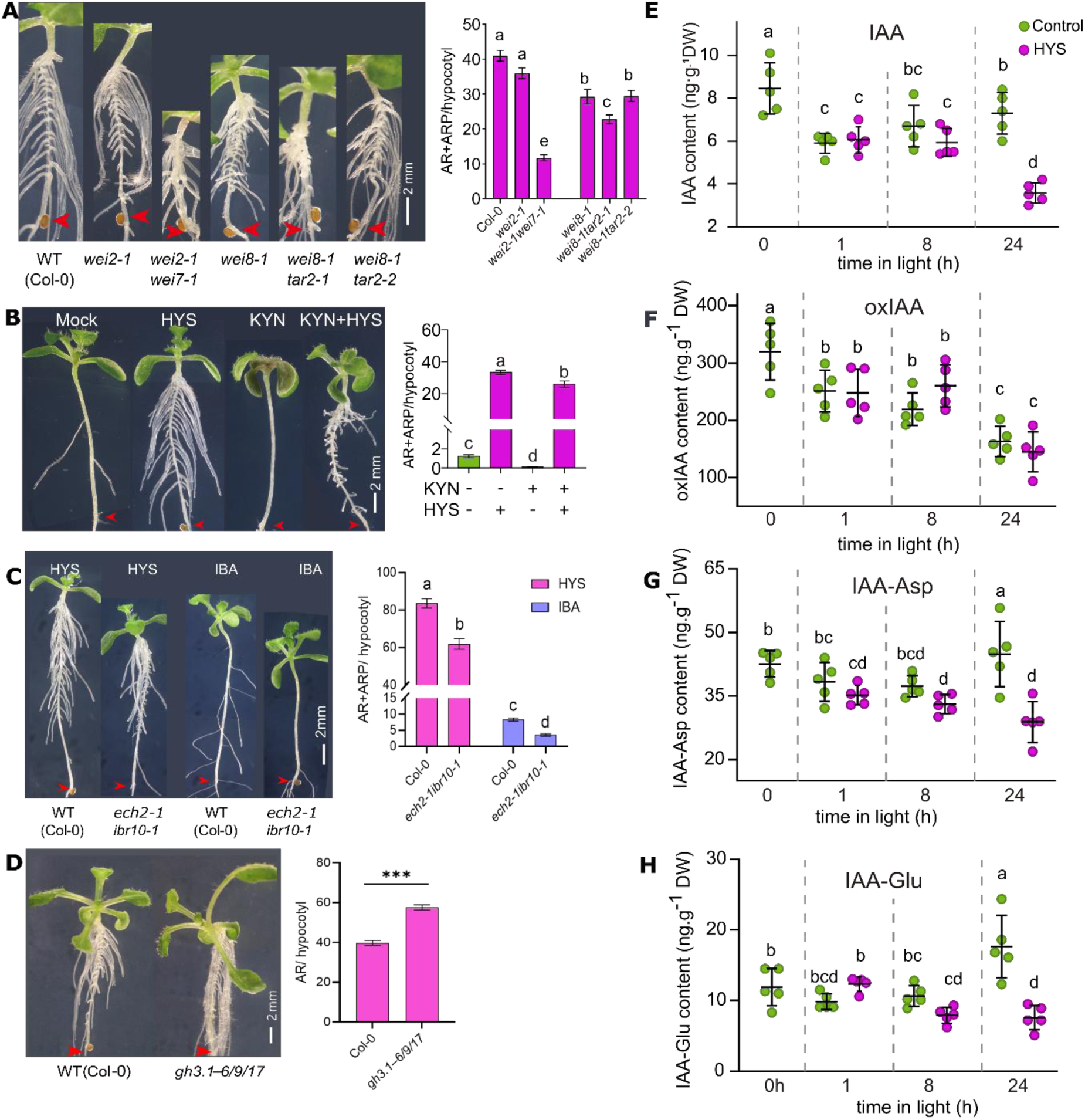
Interplay between HYS action and IAA biosynthesis. **A)** AR formation in IAA biosynthesis mutants treated with HYS (10 µM). n = 39/39/28/40/20/20, statistical significance was assessed using one-way ANOVA followed by an LSD post-hoc test. **B)** AR formation in WT treated with 10 µM HYS in the presence or absence of KYN (200 µM). **C-D)** AR formation in the *ech2-1 ibr10-1* double mutant treated with HYS (10 µM) or IBA (10 µM) n = 20/20/15/25, and in the *gh3.1–gh3.6 gh3.9 gh3.17* octuple mutant treated with HYS (10 µM) n = 30 and 35. For the *ech2-1 ibr10-1* experiment, statistical significance was assessed using two-way ANOVA followed by Tukey’s multiple-comparison test, and different letters indicate significant differences among groups (*P* < 0.05). For the *gh3.1–gh3.6 gh3.9 gh3.17* experiment, statistical significance was assessed using a two-tailed unpaired Student’s *t*-test, and an asterisk indicates a significant difference between genotypes (*P* < 0.05). **E-H).** Evolution of the content of IAA, oxIAA, IAA-Asp and IAA-Glu during deetiolation in the presence or absence of HYS (10 µM). n=5. statistical significance was assessed using one-way ANOVA followed by an LSD post hoc test, Lettering indicates statistical groups (*P* < 0.05).

**Supplementary Fig S1.**
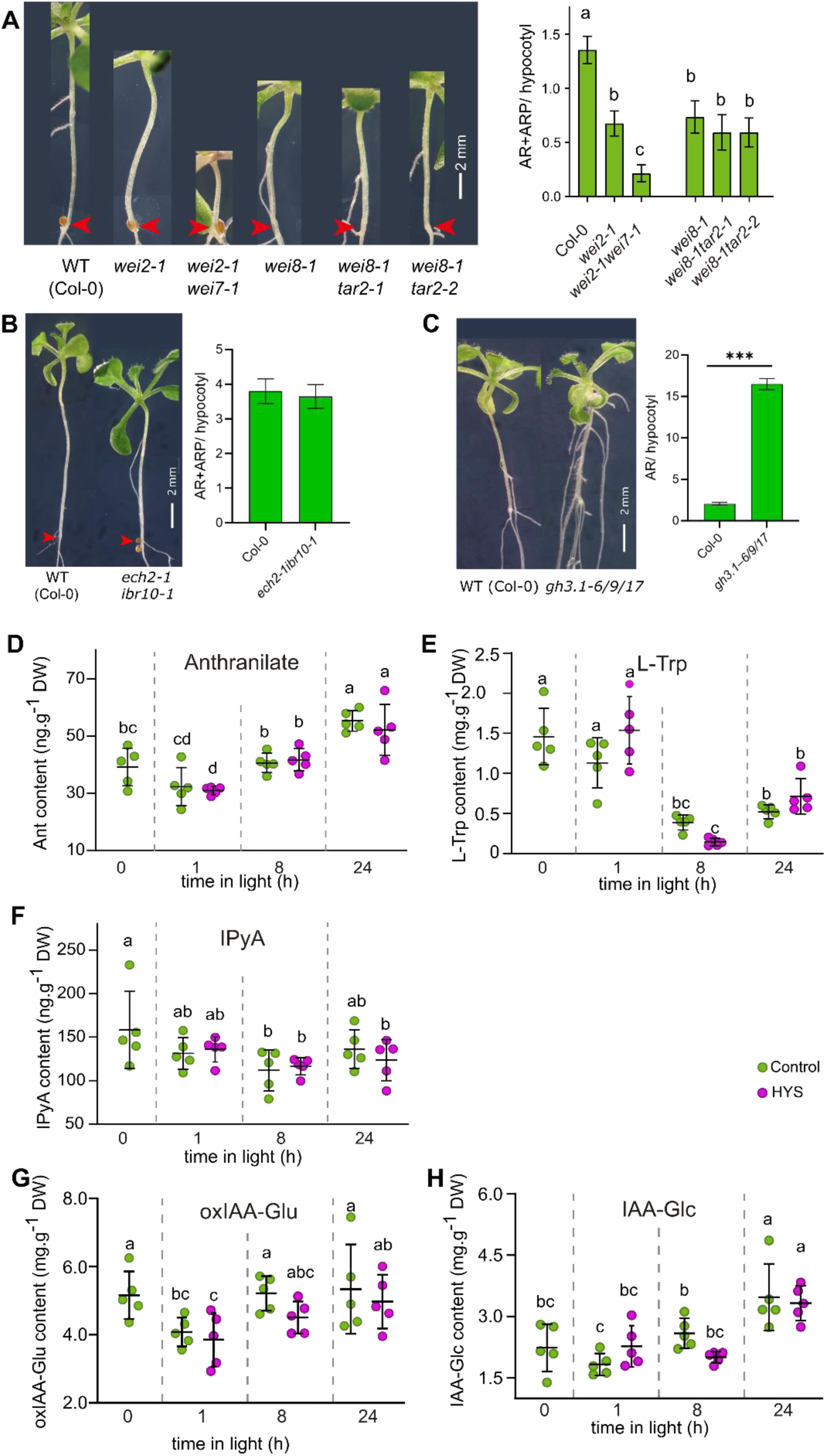
**A)** AR formation in IAA biosynthesis mutants treated for 10 days with DMSO after etiolation. n= 37/40/28/40/20/22, statistical significance was assessed using one-way ANOVA followed by an LSD post-hoc test, **B)** AR formation in the *ech2-1 ibr10-1* double mutant treated for 10 days with 0.1 % DMSO after etiolation. n = 15 and 20, and **C)** in the *gh3.1–gh3.6 gh3.9 gh3.17* octuple mutant treated for 10 day with 0.1 % DMSO. n = 35 and 35. statistical significance was assessed using a two-tailed unpaired Student’s *t*-test, and an asterisk indicates a significant difference between genotypes (*P* < 0.05). **D-H).** Evolution of the content of Anthranilate **(D)**, L-Trp **(E)**, IpyA **(F)**, oxIAA-Glu **(G),** and IAA-Glc **(H)** in seedlings during deetiolation in the presence or absence of 10 μM HYS. n=5. statistical significance was assessed using one-way ANOVA followed by an LSD post-hoc test, Lettering indicates statistical groups (*P* < 0.05).

To determine if HYS would induce AR through effects on IAA metabolism, we analyzed the impact of HYS on IAA and IAA metabolites (Figs. 1E-H; Supplementary Fig. S1D-H). HYS had no significant effects on the metabolite profiles within 8 h of treatment. After 24h, IAA and its amino acid conjugates IAA-Asp and IAA-Glu were significantly reduced by HYS treatment. This negative effect of HYS on free IAA content precludes that HYS induces AR by stimulating IAA biosynthesis or inhibition of IAA inactivation pathways.

### Chemical HYS instability does not account for HYS activity

Given that endogenous IAA metabolism alone could not account for HYS activity, we next asked whether HYS itself could be converted into a bioactive auxin or auxin-like molecule. Structurally, HYS contains a 4-chloro-2-methylphenoxyacetic acid (MCPA) moiety, derived from the synthetic auxin MCPA, linked through an amide bond to a 4-iodo-2-methylaniline moiety. As a first possibility, we examined whether HYS undergoes spontaneous chemical conversion into bioactive derivatives. Liquid chromatography-mass spectrometry (LC-MS) of a HYS solution in DMSO exposed to light in our plant incubators for 5 days identified two additional peaks that correspond to: (1) 2-(4-chloro-2-methylphenoxy)-*N*-(o-tolyl)acetamide (hereafter HA8), a HYS analogue lacking the iodine substituent (∼8.12%) and (2) a methylsulfenyl HYS derivative (∼1.28 %) (Supplementary Fig. S2A-B). This indicates that any of these derivatives could define the active molecule through which HYS induces AR.

To determine whether this major derivative retains auxin activity, we examined auxin responses using the synthetic auxin reporter DR5::GUS. HA8 induced a stronger DR5::GUS response in the hypocotyl than HYS, indicating that deiodination enhances the apparent auxin activity of the molecule and supporting the possibility that HA8 formation could contribute to HYS bioactivity (Supplementary Fig. S3). However, the low amount of produced second derivative seems unlikely to contribute significantly to HYS activity. To test its contribution, we dissolved HYS in acetonitrile to avoid its production. No significant difference in HYS-induced AR formation was apparent when DMSO or acetonitrile were used as HYS solvents (Supplementary Fig. S2C). Together, this suggests that among the detected chemical HYS derivatives, HA8 is the most likely to contribute to or modify HYS activity.

**Supplementary Figure S2.**
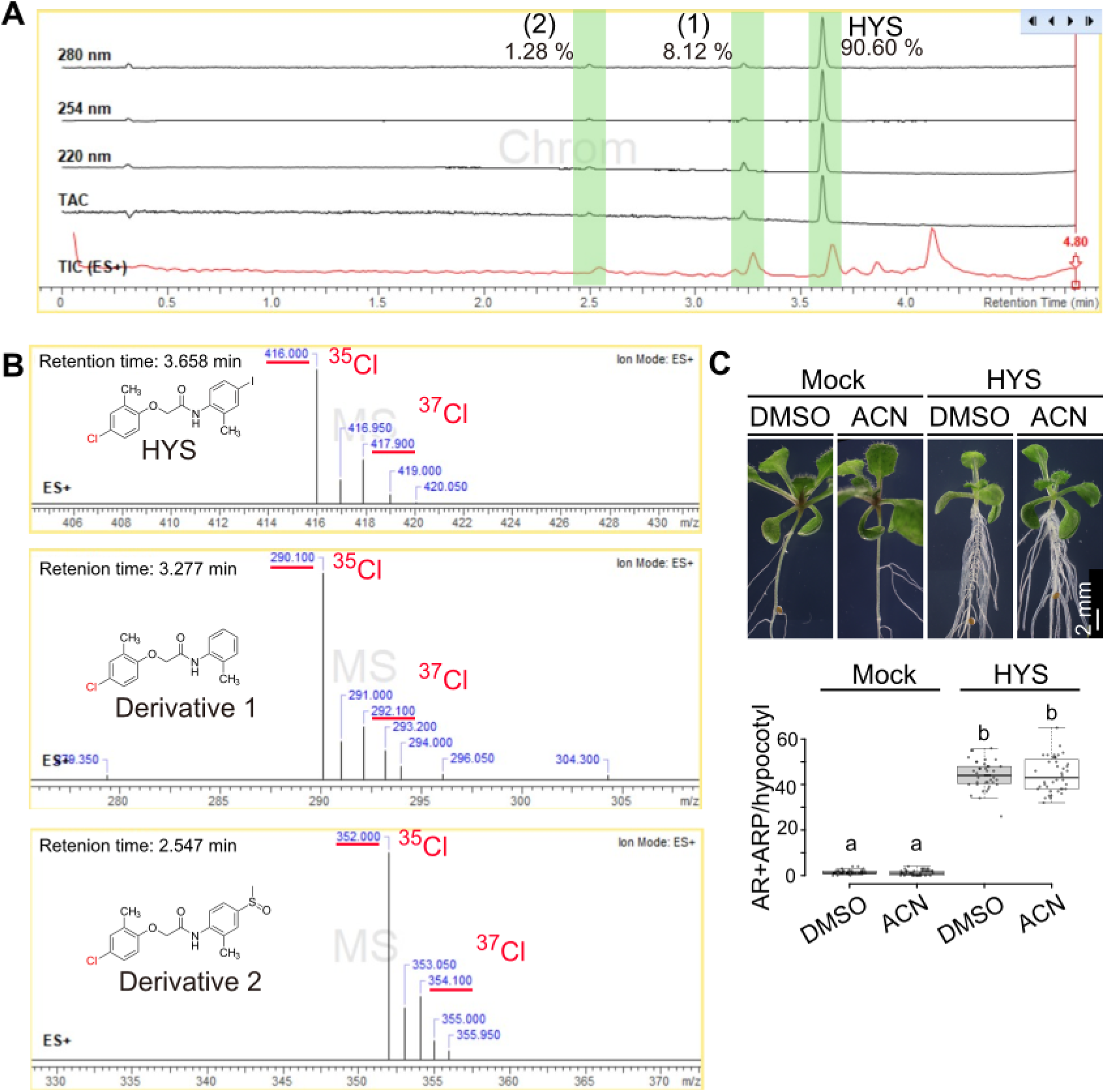
Analysis of light-induced chemical instability of HYS. **A)** LC-MS **c**hromatogram of a solution of HYS dissolved in DMSO that was exposed to light in the plant growth room for 5 days. Green bars highlight peaks of HYS (RT: 3.658 min), Derivative 1 (1) (RT: 3.277 min), and Derivative 2 (2) (RT: 2.547 min). **B)** Mass spectra of HYS, Derivative 1 and Derivative 2. The presence of m/z variants that differ 2 Da in a ratio of ∼1:3 is typical for the presence of Cl that is present in the MCPA moiety, based on the natural abundance of ^35^Cl and ^37^Cl isotopes. **C)** Adventitious root induction by HYS dissolved in DMSO vs acetonitrile. n = 40; One-way ANOVA in combination with Tukey’s multiple comparisons test, significant differences (*P* ≤ 0.05) are indicated by different lowercase letters. Central bands in the box plots show the medians; box limits indicate the 25^th^ and 75^th^ percentiles as determined by R software; whiskers extend 1.5 times the interquartile range from the 25^th^ and 75^th^ percentiles, outliers are represented by dots.

**Supplementary Figure S3.**
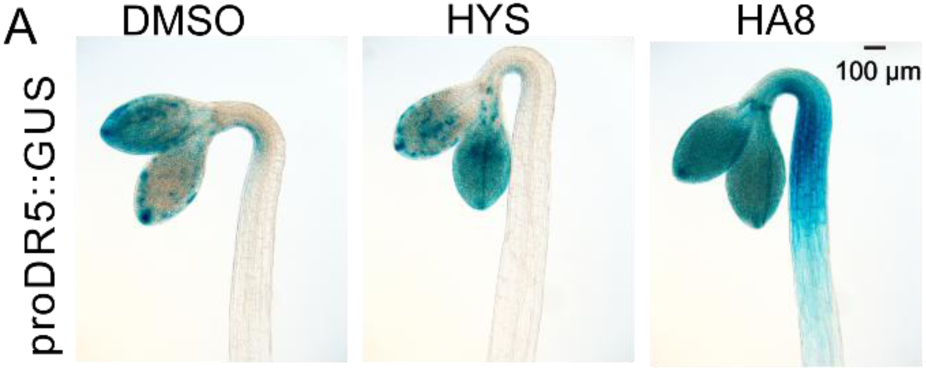
HYS, HA8, induce distinct auxin-response dynamics. **A)** DR5::GUS activity in the apical hook and hypocytl of etiolated seedlings after 3 h of 0.1 % DMSO, 10 μM HYS, 10 μM HA8.

### HYS is a proauxin that releases the synthetic auxin MCPA

Next, we explored the relevance of this light-induced conversion to HA8 during HYS-induced AR formation. For contributing to HYS activity, this molecule should be detectable at a timepoint that HYS elicits auxin responses. Therefore, we performed high-Resolution LC-MS on seedlings exposed to 10 μM HYS for 24 h under light conditions (Fig. 2A). HA8 was not detectable in the seedlings nor in the corresponding media, suggesting that light-induced HA8 production is too low to contribute to the early HYS-induced auxin responses. Instead, samples contained detectable quantities of MCPA, a synthetic auxin and its hexosylated conjugate (MCPA-hex). The hexosylated MCPA conjugate likely represents a downstream detoxification product formed by glycosylation (Bärenwald et al., 1993; Torra et al., 2024). Seedlings treated with 10 μM HA8 for 24h also contained MCPA and MCPA-hex (Fig. 2B). The absence of MCPA or MCPA-hex in either growth media indicates that they were generated *in planta* (Supplementary Fig. S4A). These results identify HYS and HA8 as proauxins that can be converted into MCPA, with the activation of HYS hydrolysis not depending on a light-induced HA8 intermediate.

**Figure 2.**
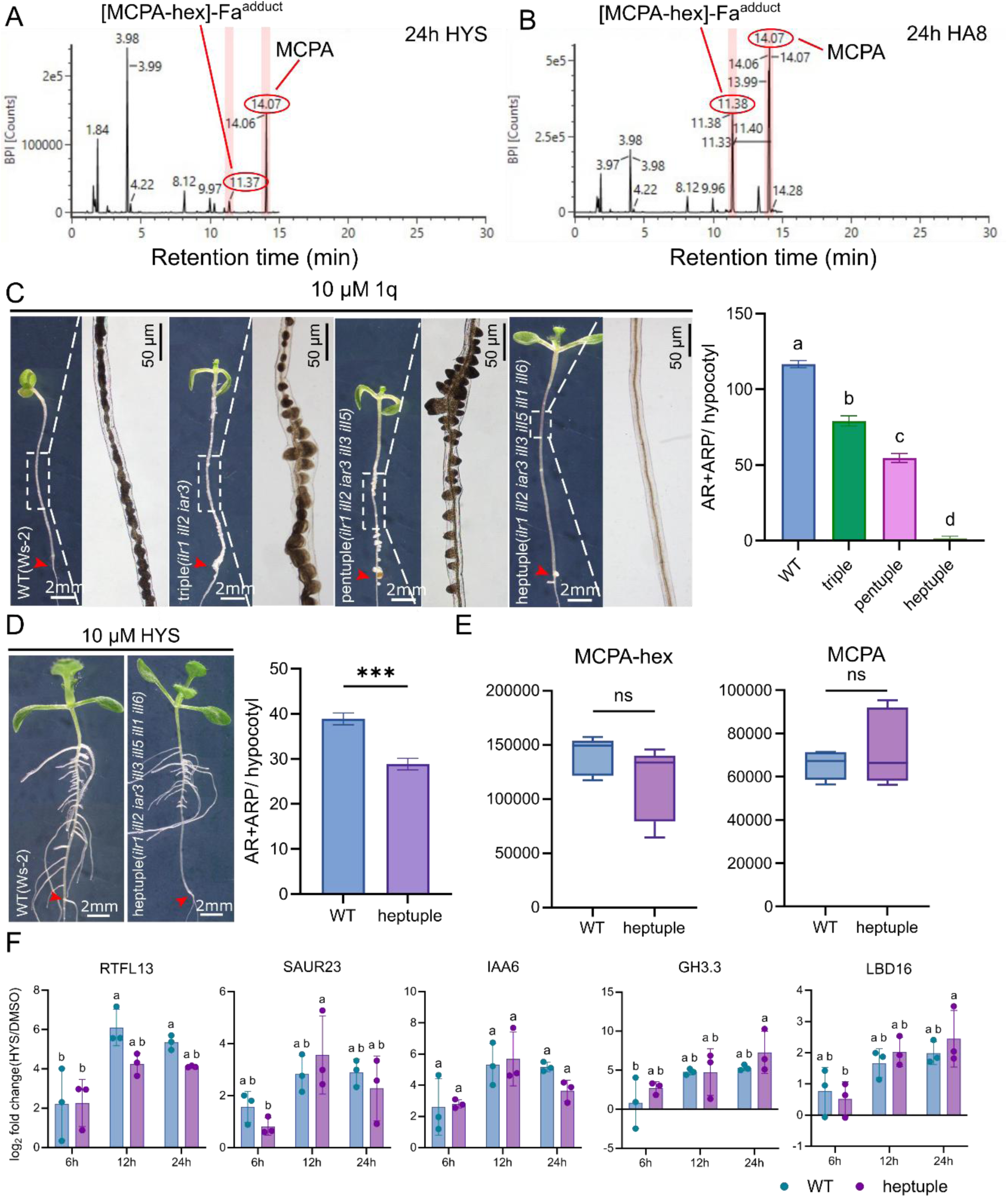
HYSPARIN is an ILR1/ILL-independent proauxin. **A-B)** LC-MS detection of MCPA and MCPA-hexose in shoots of etiolated seedlings treated with 10 μM HYS (A), or HA8 (B) for 24h in light. MCPA-hex was detected with a formic acid adduct (MCPA-hex-FA). **C**) AR formation in *ilr1-1 ill2-1 iar3-2*, *ilr1-1 ill2-1 iar3-2 ill3-1 ill5-1*, and heptuple mutants after 10-day treatment with 10 µM 1q following etiolation. n = 30/27/22/31. Statistical analysis was performed using one-way ANOVA followed by LSD post-hoc test (different letters indicate significant differences, *P* < 0.05) **D)** AR formation in heptuple mutants treated with 10 µM HYS for 10 days after etiolation and quantification of AR and ARP per seedling. n = 41 and 43. Statistical analysis was performed unpaired Student’s *t*-test (***, P ≤ 0.001), as appropriate. **E)** Levels of MCPA and MCPA-hexose in 3-day-old etiolated WT and heptuple seedlings after48 h of treatment with 10 μM HYS. n = 5. Statistical significance was assessed using a two-tailed unpaired Student’s *t*-test. ns, not significant. **F)** qRT-PCR analysis of 3-day-old etiolated wild type (Ws-2) and heptuple seedlings treated with 0.1% DMSO or 10 µM HYS for 6, 12, and 24 h. Expression levels were normalized to *UBQ10*. Data are presented as means ± SD (n = 3 biological replicates). Statistical significance was assessed by two-way ANOVA followed by Tukey’s multiple comparisons test. Different letters indicate statistically significant differences between groups (*P* < 0.05), whereas groups sharing the same letter are not significantly different.

**Supplementary Figure S4.**
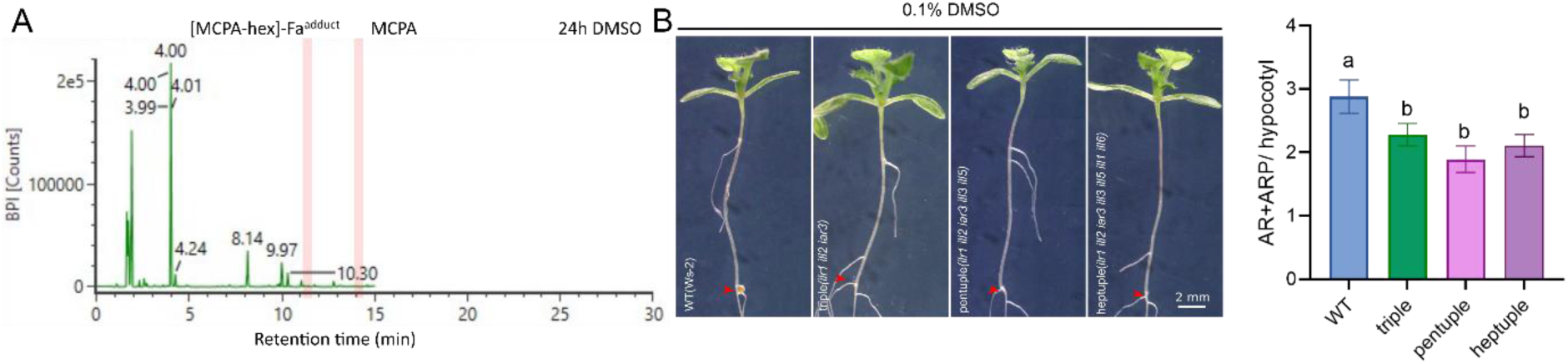
Control LC–MS analysis and AR phenotypes of *ilr/ill* mutants. **A)** LC–MS detection of MCPA and MCPA-hexose in shoots of etiolated seedlings treated with 0.1% DMSO for 24 h under light conditions. Pink boxes indicate the expected positions of MCPA and MCPA-hexose. Neither MCPA nor MCPA-hexose was detected at the corresponding positions in the DMSO-treated samples. **B)** AR formation in *ilr1-1 ill2-1 iar3-2*, *ilr1-1 ill2-1 iar3-2 ill3-1 ill5-1*, and heptuple mutants after 10 days of treatment with 0.1 % DMSO following etiolation. n = 25/25/28/28. Statistical analysis was performed using one-way ANOVA followed by an LSD post-hoc test. Different letters indicate significant differences (*P* < 0.05).

**Supplementary Figure S5.**
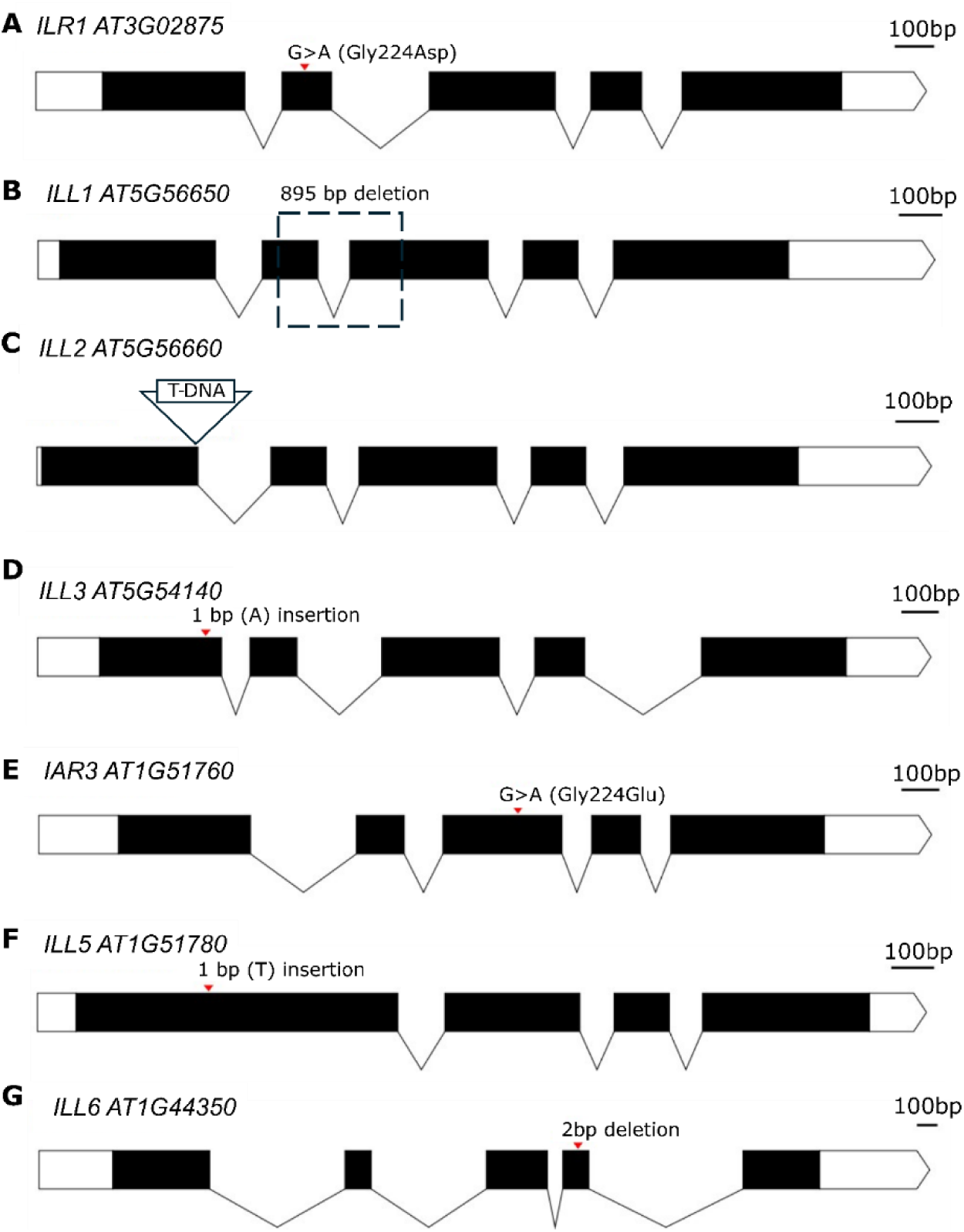
Gene structures and mutation sites of the *ILR1/ILL* heptuple mutant. Gene structures and mutation sites of (A) *ilr1*, (B) *ill1*, (C) *ill2*, (D) *ill3*, (E) *iar3*, (F) *ill5*, and (G) *ill6* alleles.

### HYS hydrolysis defines a novel proauxin activation pathway

The conversion of HYS to MCPA requires cleavage of the amide bond linking the MCPA and 4-iodo-2-methylaniline moieties. We sought to investigate the enzymes responsible for this hydrolysis. Members of the ILR1/ILL family have been implicated in the hydrolysis of the amide bond in the bipartite proauxins 4-chlorophenoxyacetic acid-L-tryptophan-OMe (named 1q) (Roth et al., 2024) and 2,4-dichlorophenoxyacetyl-L-aspartic acid and 2,4-dichlorophenoxyacetyl-glutamic acid (Chiu et al., 2018). To explore if ILR1/ILL could also hydrolyse the amide bond in HYS, we evaluated direct enzymatic hydrolysis of HYS using a bacterial enzyme assay. Clarified cell lysates from AtILL2-, AtILL6-, AtILR1-, or AtIAR3-producing bacterial cultures was incubated with HYS. The canonical ILR1/ILL substrate IAA-Ala was efficiently converted to free IAA. In contrast, no MCPA could be detected in any HYS-treated lysate under identical conditions (Supplementary Table S1). This indicates that the tested ILR1/ILLs do not directly hydrolyse HYS to MCPA.

To account for potential intermediate metabolic steps or co-factors not captured by the bacterial system, we next employed a genetic approach. To account for functional redundancy, we generated a higher-order *ilr1/ill* mutant that contains inactivating mutations in all seven *ILR1/ILL* genes (hereafter heptuple), by combining previously produced *ilr1/ill* higher order mutants (Roth et al., 2024). Consistently with redundancy in 1q hydrolysis between the different ILR1/ILLs, the heptuple was more resistant to 1q than the *ilr1 ill2 iar3* triple mutant and the *ilr1 ill2 iar3 ill3 ill5* pentuple mutant (Fig. 2C, Supplementary Fig. S4B and S5A-G). Despite the very strong 1q resistance in the heptuple, the mutant still formed many AR in response to HYS, albeit to significantly lower numbers (Fig. 2D). This indicates a role for ILR1/ILL enzymes in HYS-induced AR formation.

To assess if this effect was at the level of HYS hydrolysis, we quantified the levels of MCPA, MCPA-hexose in HYS treated heptuple mutants. In contrast to the significant AR-defect, the levels of MCPA and MCPA-hexose in the heptuple mutant were not significantly different from WT (Fig. 2E), corroborating the *in vitro* results that ILR1/ILL are not involved in the hydrolysis of HYS. Moreover, the dynamics of HYS-induced auxin responsive genes were not changed in the heptuple (Fig. 2F). Jointly, these data show that the enzymatic hydrolysis of the shoot-specific proauxin is independent of ILR1/ILL amidohydrolases.

### Structure activity relationship analysis of HYS bioactivity

Having established that HYS is an MCPA-releasing proauxin whose hydrolysis is independent of ILR1/ILL amidohydrolases, we next asked which structural features determine its characteristic for its AR-promoting activity. Structurally, HYS is a bipartite molecule in which the synthetic auxin MCPA is linked via an amide bond to 4-iodo-2-methylaniline (Fig 3A). To define the structural determinants of HYS activity, we generated a series of HYS analogues (HA) carrying systematic deletions or substitutions of the different side groups (Fig. 3B,C, Supplementary Fig. S6,C). Each analogue was applied to etiolated seedlings and assessed for effects on adventitious root (AR) and lateral root (LR) formation as well as primary root (PR) growth. As controls we included DMSO, HYS, and two synthetic auxins, NAA and MCPA. As expected, both auxins induced non-emerging AR primordia and LR primordia, and inhibited root growth and severely impaired shoot development. In contrast, HYS preferentially stimulated the development of emerged AR, while having only limited effects on LR development and PR growth (Fig. 3B). The phenotypic profile of each HA was used to group them relative to DMSO, HYS, or the synthetic auxins (Fig. 3C,D).

**Figure 3.**
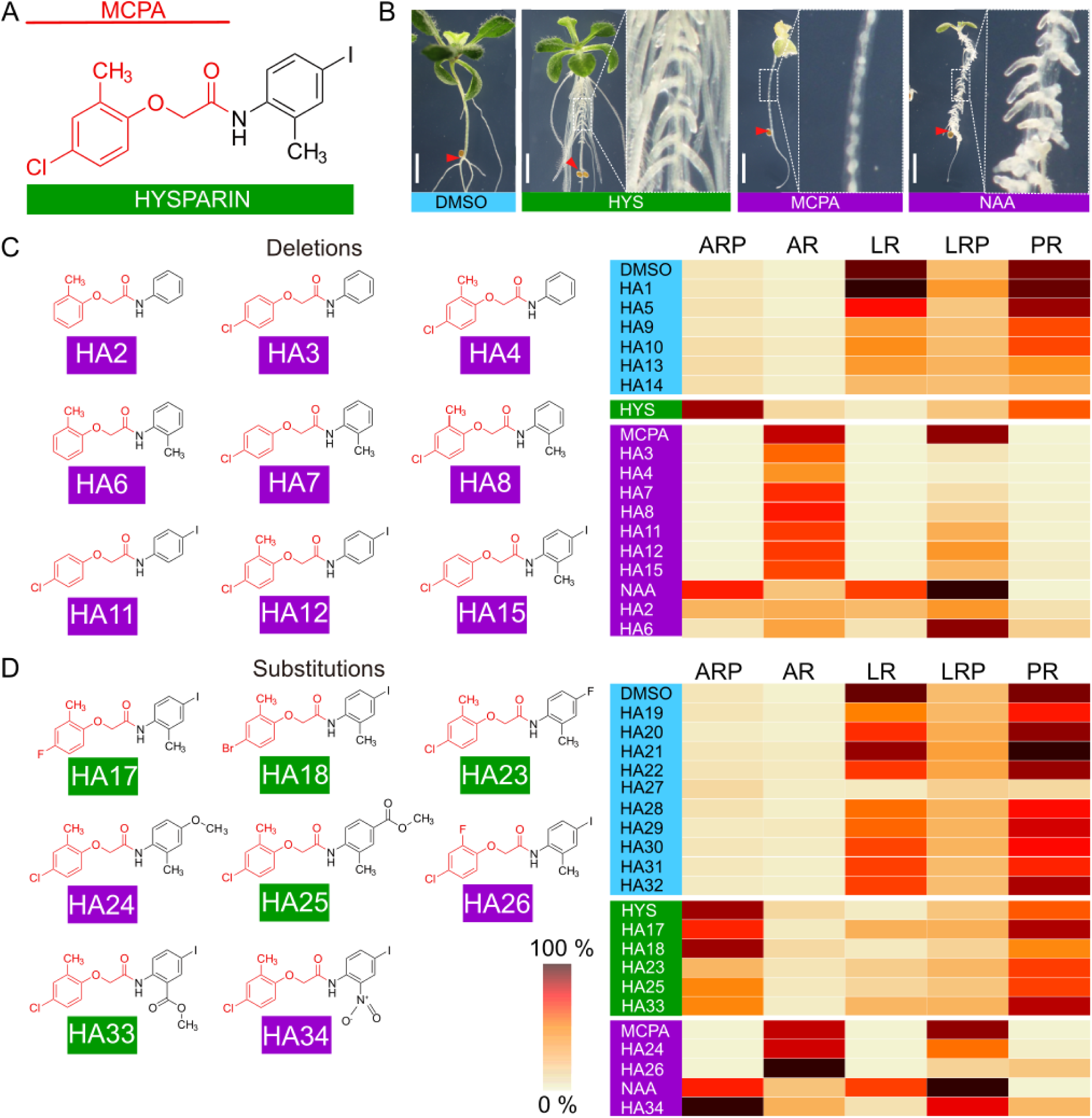
Structure Activity Relationship of HYSPARIN. **A)** Chemical structure of HYSPARIN (HYS) with indication of the MCPA-like moiety. **B)** Etiolated seedlings treated in light with DMSO (0.1 %), HYS, MCPA and NAA. **C)** Side-chain deletion analysis of HYS activity. Left, structures of HYS analogs (HA) with auxin-like bioactivity. Right, summary of phenotypic effects of HA1-HA15 relative to DMSO, HYS and MCPA/NAA. **D)** Side-chain substitution analysis of HYS activity. Left, structures of HYS analogs (HA) with HYS-like or auxin-like bioactivity. Right, summary of phenotypic effects of HA17-HA14 relative to DMSO, HYS and MCPA/NAA. All treatments were done on etiolated seedlings treated for 10 day in light with indicated compounds (10 μM). Scale is 2 mm. ARP = non-emerged adventitious root primordia; AR = emerged adventitious roots; LRP = non-emerged lateral root primordia; LR = emerged lateral roots; PR = primary root length. All values were scaled to the maximum value per phenotypic class and displayed using a color-coded scale.

First, we analyzed 15 HYS-analogues in which one or more substituents were removed from the HYS scaffold (HA1-HA15) (Fig. 3C; Supplementary Fig. S6A). Remarkably, each side chain deletion severely modified HA activity (Fig. 3C; Supplementary Fig. S6B), indicating that the structure of HYS closely approaches the minimal structural requirements for its distinctive, AR-selective bioactivity. Six deletion analogues grouped phenotypically with the DMSO control, indicating that the corresponding deletions largely abolished activity. Notably, each of these inactive HAs lacked the 4-chloro substituent of the MCPA moiety, indicating its central importance for bioactivity. In contrast, the remaining deletion HAs induced many non-emerging AR and LR primordia, and strongly inhibited primary root growth, resembling the phenotypic effects of MCPA and NAA (Fig. 3C; Supplementary Fig. 6B). This suggests that the deleted side groups within HYS suppress a broad auxin-like activity and thereby contribute to HYS selectivity.

Because MCPA belongs to the chlorinated phenoxyacetic acid class of synthetic auxins, we hypothesized that the MCPA moiety is primarily responsible for the auxin-like activities to HYS. If so, modifications at positions known to affect the activity of chlorinated phenoxyacetic acids should predictably affect HYS bioactivity.

Consistent with this expectation, substitution of the 4-chloro group with alternative halogens (fluorine in HA17 and bromine in HA18) preserved HYS-like, AR-selective activity (Fig. 3D; Supplementary Fig. S6D). In contrast, all other substitutions at this position abolished its activity (HA19–HA22) (Fig. 3D; Supplementary Fig. S6D). Similarly, replacement of the 2-methyl group with fluorine yielded an analogue with strong auxin-like activity, whereas larger substituents at this position resulted in inactive compounds (HA27–HA32). Together, these findings support the notion that the MCPA moiety determines the auxin-like activity embedded within the HYS scaffold.

Given that free MCPA acts as a non-selective auxin, we reasoned that the 4-iodo-2-methylaniline moiety contributes to the tissue specificity of HYS. Since deletion of either the 2- or 4-substituent produced auxin-like analogues, we further explored the consequences of substitutions at these positions (Fig. 3D). Although electron-withdrawing substituents might be expected to promote amide hydrolysis, no clear relationship between predicted electronic effects and biological activity emerged from the dataset. For example, replacement of the 4-iodo group with fluorine (HA23) or a methyl ester substituent retained HYS-like activity, whereas substitution with a methoxy group generated an auxin-like analogue (HA24). Likewise, replacement of the 2-methyl group with a methyl ester yielded a HYS-like analogue (HA33), while substitution with a nitro group produced an auxin-like compound (HA34). These observations suggest that the contribution of the aniline moiety to HYS activity cannot be explained by simple electronic effects on amide stability.

Collectively, our SAR analysis demonstrates that HYS contains an auxin-active MCPA module whose activity is constrained by multiple structural features within the 4-iodo-2-methylaniline moiety to achieve the distinctive AR-selective bioactivity of HYS.

**Supplementary Figure S6.**
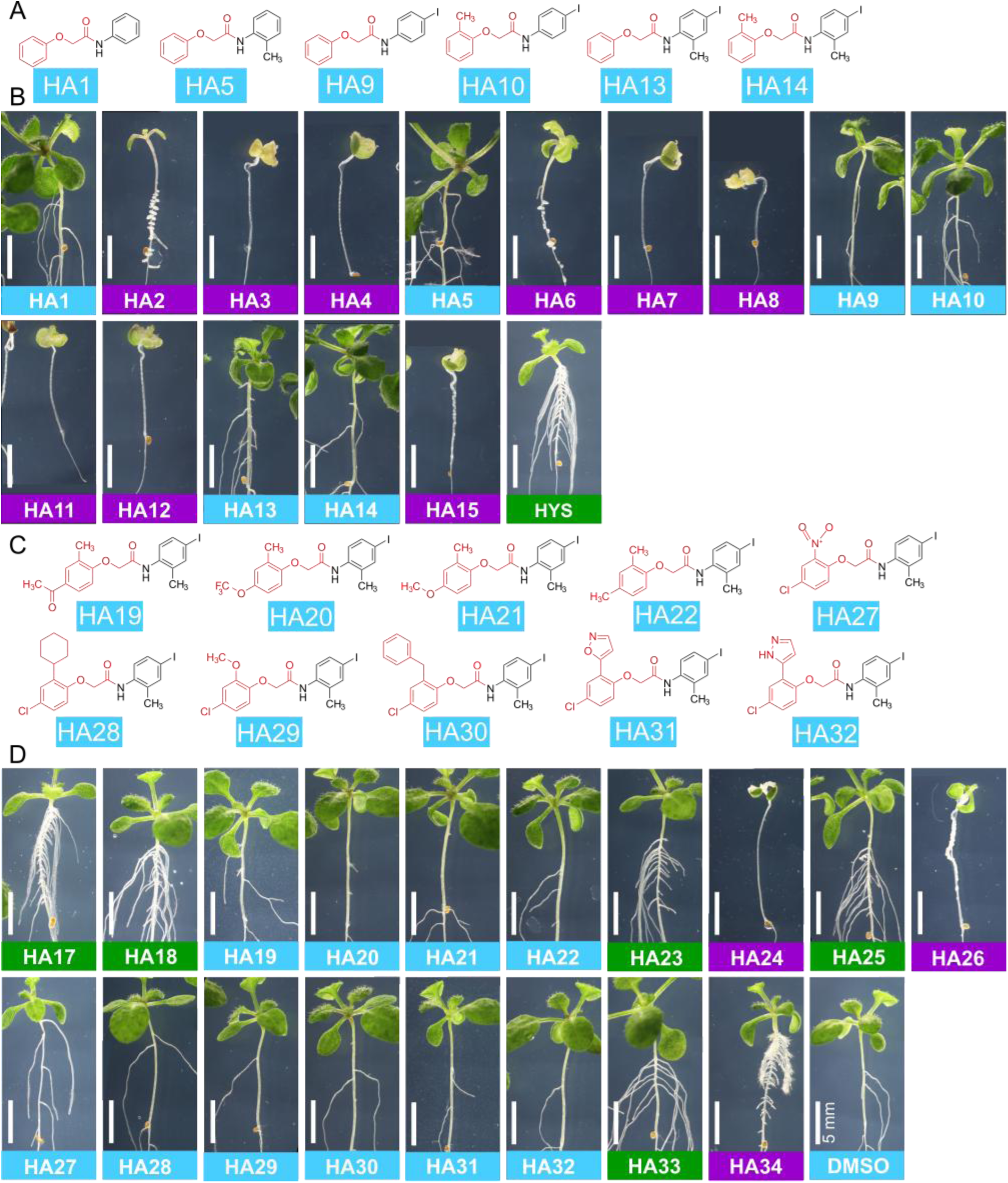
Structures of HYS analogs and their effect on AR formation. **A)** Structures of HAs from the deletion analysis with limited activity. **B)** Overview of the AR phenotype of etiolated WT treated with the deletion series of HYS analogs HA1-HA15. HYS is included as HA16 for reference. **C)** Structures of HAs from the substitution analysis with limited activity. **D)** Overview of the AR phenotype of etiolated WT treated with the substitution series of HYS analogs HA17-HA34. DMSO was included as a negative control. All HAs were applied at 10 μM for 10 days in light.

### Structural determinants of shoot-specific auxin activity

To dissect the structural determinants of HYS specificity at higher resolution, we examined the spatial activation pattern auxin responses using the synthetic auxin reporter DR5::GUS. We initially focused on HA4, HA8, and HA12, which differ from HYS only by deletions within the 4-iodo-2-methylaniline moiety while retaining the auxin moiety stable.

We compared activity of these analogs with MCPA, HYS, and the unrelated proauxin 1q (Fig. 4A-C; Supplementary Fig. S7A.). As expected, HYS selectively induced DR5::GUS activity in the hypocotyl, whereas MCPA and 1q activated auxin responses throughout the seedling. Consistently with their strong auxin-like phenotypic response in the SAR analysis, the auxin response to deletion analogs HA4, HA8, and HA12 was substantially stronger than to HYS, indicating that both the 2-methyl and 4-iodo substituents contribute to restricting the auxin activation rate of HYS. In addition to this quantitative effect, these HA variants also displayed important differences in tissue-specificity (Fig. 4D-E; Supplementary Fig. S7B). Single deletions of either the 4-iodo group or the 2-methyl group expanded the auxin response to the shoot-root junction, with limited to no induction in the root meristem. Deletion of both the 2-methyl and 4-iodo-groups led to a clear auxinic activity in the root meristem. This suggests that the 4-iodo-2-methylaniline group determines the shoot specific activity of HYS. Consistently, the spatial activity of HYS analogues that contain an intact 2-methyl,4-iodoanilide group, but contain a different auxinic moiety, remained restricted to the shoot.

**Figure 4.**
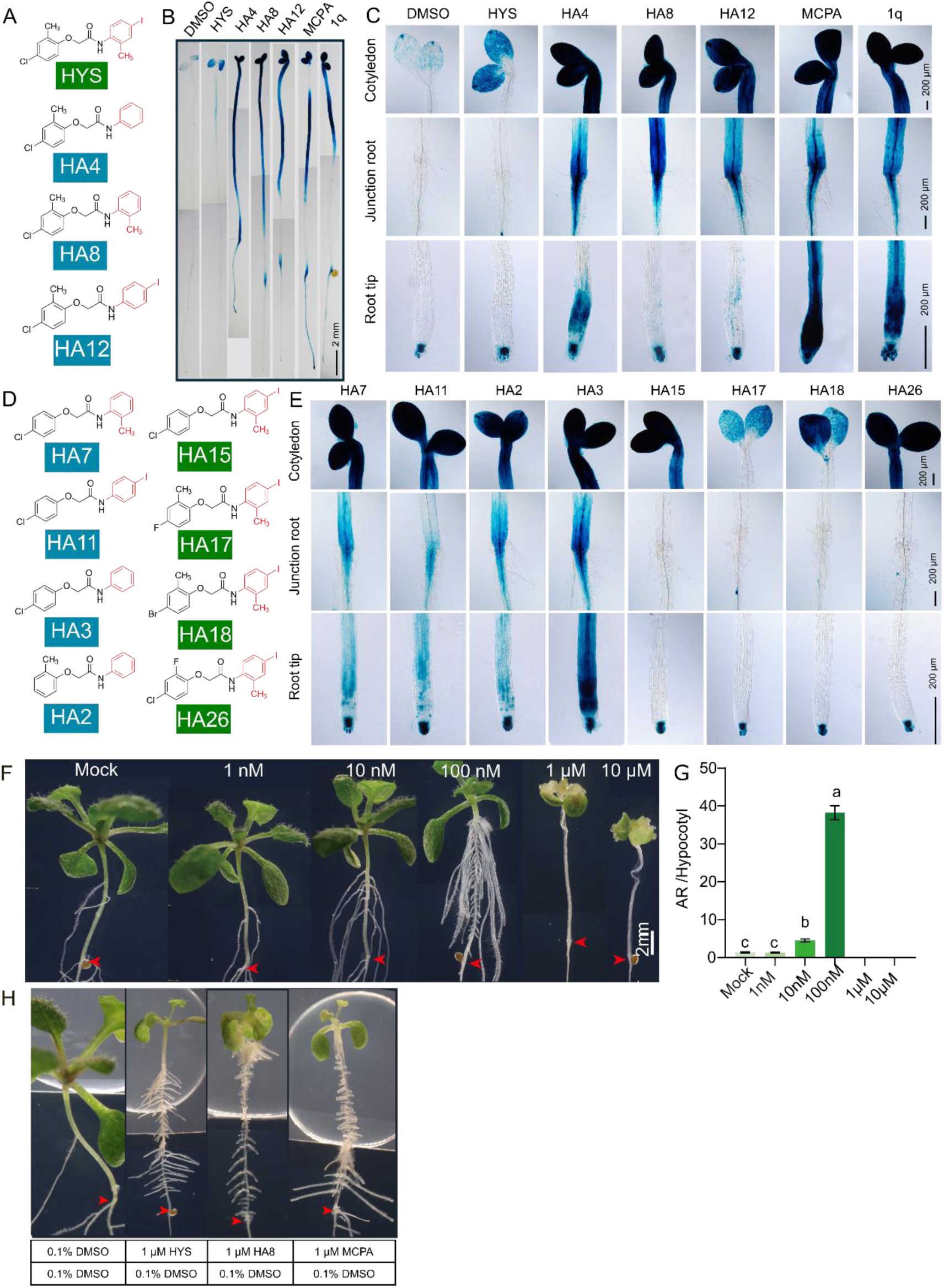
Synthetic HYS derivatives display diverse biological activities. **A) Chemical structures of HYS, HA4, HA8, and HA12. B– C)** DR5::GUS activity in (B) whole seedlings and (C) cotyledons, hypocotyl–root junctions, and root tips after treatment with 0.1% DMSO, HYS, HA4, HA8, HA12, MCPA, or 1q for 24 h. All compounds were applied at 10 μM. **D)** Chemical structures of HA2, HA3, HA7, HA11, HA15, HA17, HA18, and HA26. Analogues with modifications of the 4-iodo and/or 2-methyl substituents are indicated in blue, whereas analogues retaining an intact 4-iodo-2-methylaniline moiety are indicated in green. **E)** DR5::GUS activity in cotyledons, hypocotyl–root junctions, and root tips after treatment with the indicated HYS analogues for 24 h. **F)** Pictures of AR phenotype in etiolated seedling treated with different doses of HA8. **G)** Quantification of AR and ARP in the seedlings shown in (F). n>40, Statistical significance was assessed using one-way ANOVA followed by an LSD post hoc test, Lettering indicates statistical groups (P < 0.05). **H)** Droplet-based assay showing representative phenotypes of 10-day-old Col-0 seedlings that were etiolated for 3 days following local application of 0.1%DMSO, 1 μM.HYS, 1 μM.HA8, or 1 μM.MCPA to shoot tissues. Treatment conditions, including droplet content and plate composition, are indicated in the table.

**Supplementary Figure S7.**
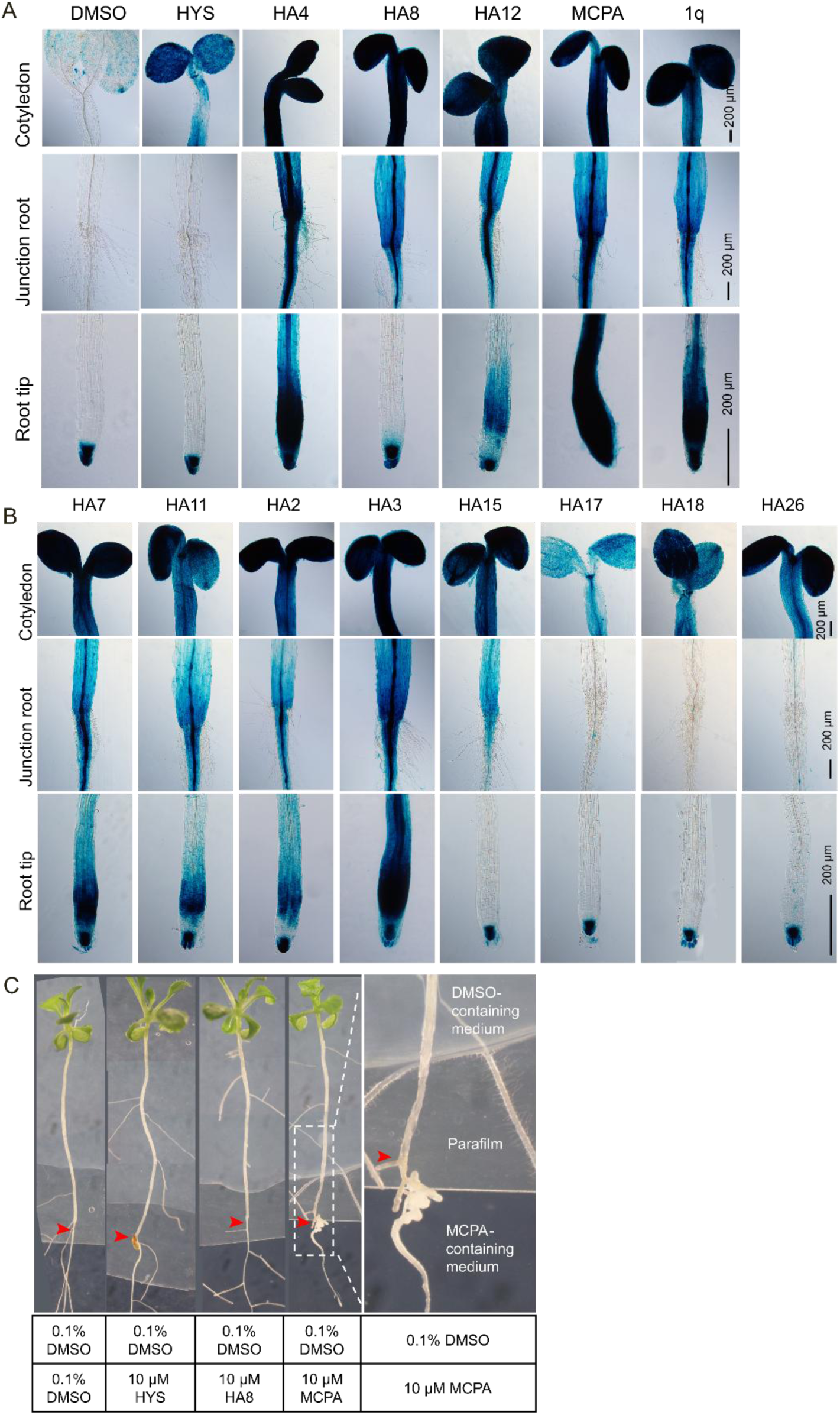
HYS and its analogues generate distinct auxin responses. **A–B)** DR5::GUS activity in cotyledons, hypocotyl–root junctions, and root tips after treatment with the indicated HYS analogues for 48 h. All compounds were applied at 10 μM. **C)** Representative phenotypes of Col-0 seedlings after 10 days of droplet-based treatment following 3 days of etiolation. Roots were exposed to medium containing 0.1 % DMSO, 10 μM HYS, 10 μM HA8, or 10 μM MCPA, while 0.1% DMSO droplets were locally applied to shoot tissues. Treatment conditions, including droplet content and plate composition, are indicated in the table.

Together, these findings indicate that the 4-iodo-2-methylaniline promoiety determines the shoot-selective delivery of phenoxy acetic acid-type auxins.

### Shoot-specific delivery of a small dose of MCPA auxin explains HYS activity

Because HA8 differs from HYS primarily in the magnitude rather than the spatial distribution of its auxin response, we next asked whether HA8 could phenocopy HYS at lower concentrations (Fig. 4F-G). Consistent with its initial characterization, HA8 displayed broad auxin-like activity at high concentrations (1– 10 μM), causing epinasty, chlorosis, and the accumulation of numerous non-emerged adventitious root primordia. Strikingly, at lower concentrations HA8 no longer interfered with shoot development or greening and instead promoted the formation of numerous well-developed adventitious roots, closely resembling the HYS phenotype. These observations suggest that the selectivity of HYS arises from its ability to generate a shoot-localized auxin response that is sufficiently strong to stimulate adventitious rooting while remaining below the threshold required to trigger broader auxin-induced developmental defects. To test this hypothesis, we selectively applied droplets of MCPA-containing medium to shoot tissues. Consistently, this shoot-application of MCPA was sufficient to induce AR in a pattern that is reminiscent of HYS activity (Fig. 4F). In contrast, root-localized application of MCPA induced an accumulation of lateral root primordia at the application site but did not promote AR formation in the hypocotyl (Supplementary Fig. 7C).

Collectively, these results show that the unique AR-selective activity of HYS derives from spatially restricted delivery of a low dose of the synthetic auxin MCPA.

## DISCUSSION

Among molecules with auxin-like activity, HYS stands out for its shoot-selective activity. It is a small molecule that combines potent induction of adventitious root formation with minimal effects on root growth and overall plant development (Zeng et al., 2023). Here, we investigated the mechanistic basis of this unique developmental selectivity. Our results support a model in which HYS functions as a shoot-selective proauxin that is metabolically converted into MCPA through an ILR1/ILL-independent pathway. This spatially restricted activation generates auxin levels that are sufficient to induce adventitious root formation while avoiding toxic, systemic auxin responses. Furthermore, our structure–activity analyses identify key substituents within the HYS scaffold that modulate both the tissue specificity and the magnitude of the released auxin signal. Together, these findings establish HYS as a proof-of-concept for achieving developmental specificity through tissue-selective prohormone activation rather than through modification of hormone perception or signaling itself. More broadly, our work highlights spatially restricted metabolic activation as a powerful strategy for uncoupling desirable hormone-regulated developmental processes from their pleiotropic consequences.

### HYSPARIN is a proauxin

Adventitious roots depends on canonical auxin signaling, but is unable to directly activate TIR1/AFB auxin receptors (Zeng et al., 2023). This indicated that HYS activates auxin signaling indirectly. The induction of IAA production or inhibition IAA turnover are unlikely mechanistic explanations, as HYS treatment reduces rather than increases endogenous IAA levels. Instead, our data support a model in which HYS functions as a proauxin that is metabolically converted into the synthetic auxin MCPA. This conclusion is supported by both the structure of HYS and its structure–activity relationships. The structure of HYS contains an MCPA moiety whose structural requirements closely mirror those governing the activity of chlorinated phenoxyacetic acid auxins (Karami et al., 2023). In particular, the 4-chloro substituent, which is essential for the activity of this auxin class, was also indispensable for HYS activity. In HYS this chlorine can be replaced by another halogen without severe impact on its AR-selective activity. Likewise, modifications at the 2-position altered the physiological profile of HYS in a manner reminiscent of the effects of analogous substitutions on chlorinated phenoxyacetic acids (Karami et al., 2023). HYS analogues lacking the 2-methyl group, or carrying alternative substituents at this position, displayed stronger and less selective auxin-like activities. Together, these observations indicate that the auxin activity of HYS resides in an MCPA-derived metabolite and that the remaining structural elements of the molecule primarily modulate the spatial and quantitative delivery of this activity.

Light exposure resulted in the slow deiodination of HYS producing the highly bioactive HYS analogue HA8. This HA8 is unlikely to act as an intermediate in HYS activation, as it did not accumulate in the plant to detectable levels within 48 h, a time points at which in the first cell divisions of HYS-induced adventitious root could be detected (Zeng et al., 2023). Instead, HYS-treated seedlings accumulated free MCPA and a hexosylated MCPA conjugate that likely reflects downstream detoxification (Bärenwald et al., 1993; Torra et al., 2024). This supports a model in which HYS releases MCPA by hydrolysis of its amide bond. These findings identify HYS as a novel proauxin (Christian et al., 2008; Savaldi-Goldstein et al., 2008; Kerchev et al., 2015; Roth et al., 2024). Uncoverying the mechanism responsible for HYS activation will therefore be an important future steps toward full understanding HYS bioactivity.

### Mechanism for shoot-specificity

Unlike conventional synthetic auxins, HYS displays a remarkable tissue-selectivity, indicating spatially restricted activation. This spatially restricted activation allows to uncouple auxin-dependent root initiation from the growth inhibition and developmental abnormalities typically associated with systemic auxin exposure. Our data suggest that the 2-methyl group in 4-iodo,2-methylanilide moiety acts as a selectivity filter impacting on shoot selective hydrolysis. Deletion series of this substituent abolished HYS shoot-specificity, activating strong auxin responses throughout the seedling.

One possible explanation is that the 2-methyl group restricts HYS hydrolysis to a limited subset of amidases. Consistent with this hypothesis, plant aryl-acylamidases involved in propanil metabolism exhibit strong sensitivity to substitution patterns on the aniline ring, and substrates carrying substituents adjacent to the amide bond are often hydrolysed less efficiently than closely related analogues (Villarreal et al., 1994). The 2-methyl group may therefore act as a structural filter that limits HYS activation to enzymes with a suitably configured substrate-binding pocket. Such a mechanism would provide a simple explanation for the shoot-selective activation of HYS and the broader activity spectrum of analogues lacking this substituent.

The activation mechanism of proauxins has thus far been elucidated for the 2,4-D-Asp and 2,4-D-Glu conjugates {Chiu, 2018 #1543}, and the proauxin 1q {Roth, 2024 #2}. These amino acid conjugated synthetic auxins structurally resemble endogenous auxin- and jasmonate-amino acid conjugates. Consistent with this similarity, these synthetic auxin conjugates are activated by members of the ILR1/ILL family of amidohydrolases (Roth et al., 2024). The ability of these enzymes to process both endogenous hormone conjugates and synthetic proauxins suggested that they may possess considerable substrate promiscuity. Surprisingly, however, HYS activation was entirely unaffected in the heptuple ILR1/ILL mutant lacking all known family members. Together with the inability of recombinant ILR1, ILL2, ILL6, and IAR3 proteins to release MCPA from HYS *in vitro*, these findings demonstrate that HYS is activated through an ILR1/ILL-independent pathway. HYS therefore reveals the existence of at least two mechanistically distinct routes for proauxin activation in plants. The ILR1/ILL-dependent one being active on amino acid conjugates such as 1q, while the ILR1/ILL-independent pathway for HYS hydrolysis targets auxins conjugated to aromatic anilides. Shoot-specific activity of the corresponding hydrolytic enzyme could thus explain the shoot-specificity of HYS. An alternative explanation could be that the shoot-specificity derives rather from shoot-selective uptake followed by activation by a ubiquitously active enzyme.

### Slow auxin delivery to the shoot for AR organogenesis

Restricted hydrolysis of HYS likely underlies its ability to promote adventitious root (AR) formation while preserving normal primordium development and overall root patterning. The formation of a complex three-dimensional organ such as an adventitious root requires tightly coordinated rounds of cell division and differentiation to establish a functional root meristem (Stoeckle et al., 2018). This process depends on a strong initial auxin input to initiate AR formation, followed by dynamically evolving auxin gradients that guide patterning and tissue organization (Motte et al., 2019). These gradients arise from the interplay between auxin biosynthesis, transport, and inactivation, which together generate spatially and temporally refined auxin maxima (Vanneste et al., 2025).

In contrast, many synthetic auxins behave as stable, freely diffusible compounds that are poorly substrates for endogenous auxin homeostasis mechanisms (Hayashi, 2021). This can lead to prolonged and spatially uniform signaling, which disrupts gradient formation and compromises organ patterning. This is evident in the strong phytotoxic effects of high concentrations of MCPA (Fig. 3B), which inhibit proper AR primordium organization and suppress normal root development rather than promoting coordinated organogenesis.

A proauxin strategy has recently been used to partially address these limitations. For example, the compound 1q functions as a cell-permeable auxin precursor that is hydrolysed intracellularly to release the active auxin 4-CPA (Roth et al., 2024). When applied in short pulses, this system improves rooting efficiency across multiple species. However, prolonged exposure to 1q or its active metabolite still leads to growth inhibition and aberrant primordium patterning (Fig. 2C), highlighting that sustained, systemic auxin release remains detrimental even when delivered as a proauxin.

In this context, HYS represents a distinct mode of auxin delivery. HYS treatment induces robust adventitious root formation while maintaining well-organized primordia, even under continuous exposure with high concentrations (Zeng et al., 2023). Notably, despite the shared shoot-restricted activation of auxin responses by HYS and HA8, they elicit phenotypically very different responses. Only when HA8 doses were reduced, normal root growth was established and well-patterned adventitious root were formed, similar to HYS. These observations indicate that the developmental outcome does not only depend on the spatial restriction of auxin release, but also on its magnitude, thereby closely adhering to how endogenous auxin homeostasis mechanisms control plant development. More broadly, these findings highlight that the selective control of plant development can be achieved by engineering tissue-selective prohormones with optimized release rates.

## Conflict of interest

The authors declare no conflict of interest.

## Acknowledgements

This work was supported by the Research Foundation – Flanders (FWO), under grant agreement 3G049622 (to SV And DG); Ghent University (BOF, PDO.2024.0015.01 to R.Wang); and the China Scholarship Council (CSC; 202404910064 to QL and 201806300036 to YZ). the Israel Science Foundation (1057/21 to R.Weinstain); and the Yuri Milner 70@70 Fellowship (to OR). KL was supported by funding from the Knut and Alice Wallenberg Foundation (KAW 2016.0352 and KAW 2020.0240) and the Swedish Research Council (VR 2021-04938).

## Author contributions

DG and SV conceived the initial idea, which was further developed by SS, DO, YZ, and QL. DG and SV designed and supervised the study. QL, YZ, SS, DO, ML, RW, HKT, GG, KL, OR, RW, JM, TH, and IV performed experiments, collected data, and provided materials. QL, YZ, SS, DO, and SV analyzed and interpreted the data. SV prepared the initial draft of the manuscript. All authors contributed to the revision of the manuscript and approved the final version.

## Data availability statement

Data supporting the findings of this work are available within the paper and its Supplementary information file.

---

**Supplementary Table S2.**
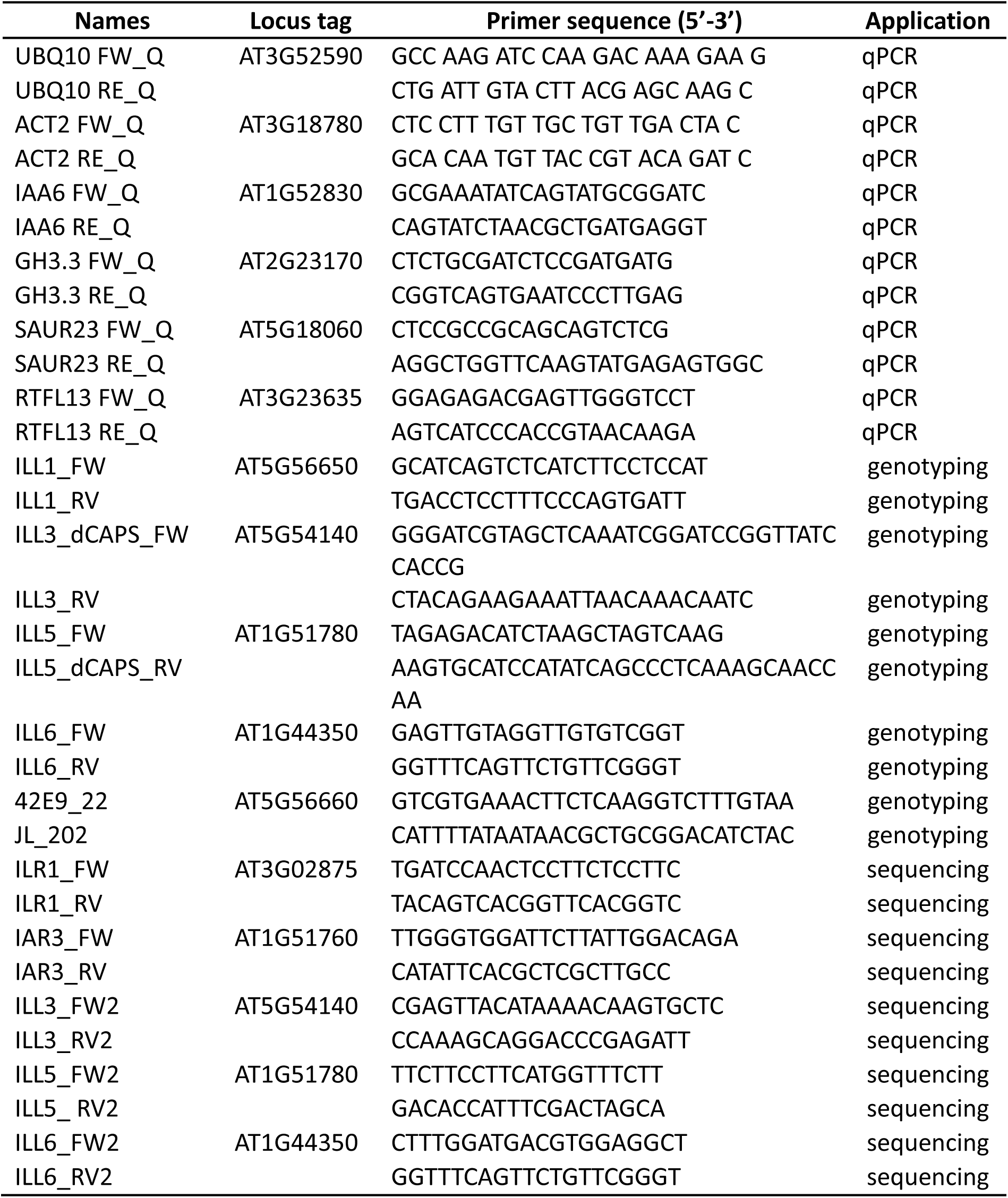
List of PCR primers used in this study.

## Materials and methods

### Plant material

The *wei2-1* (Stepanova et al., 2005), *wei2-1wei7-1* (Fattorini et al., 2017), *wei8-1* (Stepanova et al., 2008), *wei8-1tar2-1* (Stepanova et al., 2008), *wei8-1tar2-2* (Stepanova et al., 2008), *wei8-1tar1-1tar2-2* (Stepanova et al., 2008), *ech2-1ibr10-1* (Strader et al., 2011), *gh3* octuple (*gh3.1-6/9/17*) (Guo et al., 2022) mutant are in Columbia-0 (Col-0) ecotype. The *ilr1-1ill2-1iar3-2* (Rampey et al., 2004), *ilr1-1ill2-1iar3-2ill3-1ill5-1* (Roth et al., 2024), *ilr1-1ill2-1iar3-2ill1-1ill6-1* (Roth et al., 2024) are in Wassilewskija (Ws-2) background. The *ilr1/ill* heptuple mutant in Ws-2 was generated by crossing *ilr1-1 ill2-1 iar3-2 ill3-1 ill5-1* and *iar1-1 ill2-1 iar3-2 ill1-1 ill6-1* and selection by genotyping, PCR-based genotyping was performed using the primers listed in Table S2. The *ill1* deletion allele was identified by PCR, whereas *ill3* and *ill5* were genotyped by derived cleaved amplified polymorphic sequences (dCAPS) using AgeI and BsaJI, respectively. The *ill6* allele was genotyped by Eco88I digestion. Digested products were analyzed on 2.5 % agarose gels. Homozygous heptuple mutants were further validated by Sanger sequencing of the *ill3*, *ill5*, *ill6*, *iar3*, and *ilr1* amplicons. The *ill2-1* allele was genotyped as previously described (Rampey et al., 2004). (Genotyping primers are listed in Supplementary Table S2). Reporters used are DR5::GUS (Ulmasov et al., 1997).

### Growth conditions and adventitious root assay

Arabidopsis seeds were surface sterilized in chlorine gas in a closed desiccator for 3-4 hours. The sterilised seeds were sown on 0.5x Murashige and Skoog (MS) medium supplemented with 0.5 % (w/v) sucrose, 0.8 % (w/v) agar and 0.05 % (w/v) MES at pH 5.7. Following the 4 °C for 4 dark days stratification treatment, the plates were incubated in the light for 8 h before being incubated for 3 days at 22 °C in the dark for etiolation. Well elongated seedlings were transferred to chemical treatments and adventitious roots phenotype were evaluated after an additional 10 days in a growth chamber (70 % relative humidity and 22 °C) with 16 h/8 h light/dark cycles (70 µmol/m^2^s).

### Preparation and observation of cleared seedings

For light microscopy, plant material seedlings were cleared with methanol and NaOH and mounted as described (Malamy and Benfey, 1997). A BX51 microscope (Olympus, Tokyo, Japan) equipped with differential interference contrast (DIC) optics were employed to count AR and ARP.

### Droplet-based local application assay

Three-day-old etiolated Col-0 seedlings were transferred to the indicated treatment plates. For shoot treatment, seedlings were transferred to medium containing 0.1 % DMSO, and droplets containing 0.1 % DMSO, 1 μM HYS, 1 μM HA8, or 1 μM MCPA were locally applied to shoot tissues. For root treatment, seedlings were grown with their roots in contact with medium containing 0.1% DMSO, 10 μM HYS, 10 μM HA8, or 10 μM MCPA, while 0.1% DMSO droplets were locally applied to shoot tissues. After 10 days of treatment, representative seedlings were imaged using an Olympus SZX9 stereomicroscope.

### GUS staining

Histochemical GUS staining was performed as previously described (Jefferson et al., 1987), with minor modifications. Seedlings were incubated in GUS staining solution containing X-Gluc at 37°C until sufficient staining developed. Chlorophyll was subsequently cleared with lactic acid, and samples were imaged using a BX51 microscope equipped with DIC optics.

### Analysis of IAA and IAA conjugate levels

For hormone analysis, Arabidopsis seeds were sown on the Parafilm strip margin to allow roots to grow over the strip. Seeds were incubated for 4 d at 4 °C in the dark, followed by germinating in the light (22 °C, 70 µmol/m2s) for 8 h incubation before incubated in the dark to induce hypocotyl elongation as described (Zeng et al., 2021). Three-day-old etiolated seedlings were transferred to a medium with or without 10 μM HYS for 1 h, 8 h, and 24 h by lifting the Parafilm strip. Five individual samples were harvested, and tissue was homogenized in liquid nitrogen. The setting of hormone extraction and internal standards are described (López-Ráez et al., 2010). Samples were filtered through Minisart SRP4 0.45μm filters (Sartorius, Goettingen, Germany) before liquid chromatography-tandem mass spectrometry (LC-MS/MS) analysis as described (Saika et al., 2007; López-Ráez et al., 2010). For each sample, tissue was processed as previously described for free IAA or IAA conjugates (Novák et al., 2012).

### Analysis of light-induced HYS conversion via LC-MS and NMR

HYS was dissolved in DMSO at a concentration of 10 mM and exposed for 5 days to a long-day regime in the growth chamber. DMSO was removed via vacuum distillation. ^1^H-NMR analysis only detected DMSO in the distillate. A LC-MS analysis was performed on the remaining product. Using a 1200 Series LC/MSD SL equipped with a Supelco ascentis express C18 column (internal diameter 4.6 mm), a UV-DAD detector, an Agilent 1100 Series MSD SL mass spectrometer with electrospray ionisation (ESI, 4000 V, 70 eV) and with a single quadrupole detector coupled to the machine. To elute the components, a solvent mixture of acetonitrile and water in different ratios was used.

To analyse the product via ^1^H-NMR, the product was dissolved in deuterated chloroform (CDCl_3_) after sample preparation. The spectra were taken by a Bruker Avance Nanobay III NMR spectrometer with a ^1^H/BB z-gradient high-resolution probe. The ^1^H NMR was taken at 400 MHz. The software used to process and display the spectra was TOPSPIN version 3.5.4. To prepare the samples for usage, the compounds were dissolved in deuterated solvents. For quantification, the whole sample that was obtained after DMSO was analyzed by ^1^H-NMR. The sum of HYS, and both degradation products was set to 100. The signal of compound 2 was close to the detection limit, and thus an estimation.

### Analysis of in planta HYS metabolism

Three-day-old etiolated Arabidopsis seedlings were transferred to 10µM HYS medium or 0.1 % DMSO as mock control. Hypocotyls were collected in five replicates after 24 h of treatment, immediately frozen in liquid nitrogen, and stored at –80 °C until extraction. The samples (approx. 200 mg plant material fresh weight) were pulverized using a Retsch MM 400 mixer mill at a frequency of 30 Hz for 2 min after adding 2 iron beads at −70°C. The plant extracts were in 800 μl of methanol and incubated at room temperature with continuous shaking for 60 min, centrifuged for 10 min with 15 000 rpm (MAX speed) and 500 μl of supernatant was elated by SPE plate followed by 400 μl methanol washing. After purification, evaporated to dryness in Vacuum-dried and reconstituted in 100 μl 10% methanol. The samples were filtered through a 0.22 μm 96-well filter plate before HPLC-MS/MS analysis. For HYS metabolism, a 10 µl extraction solution was injected into a PR HR-LC-MS in positive and negative mode with targeted analysis for HYS, MCPA, MCPA-hex and HA8. Standards were used for HYS, MCPA and HA8.

### Gene expression analysis

Arabidopsis Ws-2 and the *ilr1/ill* heptuple mutant seedlings were grown according to the AR assay described above and treated with 10 μM HYS or 0.1% DMSO for 0, 6, 12, or 24 h. Approximately 100 mg of seedlings were collected per sample, immediately frozen in liquid nitrogen, and used for RNA extraction. Total RNA was extracted using the ReliaPrep™ RNA Tissue Miniprep System (Promega, Z6112) according to the manufacturer’s instructions. Following by reverse transcribed to cDNA using the GoScript Reverse Transcription System (Promega, A5001). Real-time quantitative PCR (RT-qPCR) was performed using the GoTaq Q-PCR Master Mix (Promega, A6001) according to the manufacturer’s instructions. Primers for UBQ1 were used as the internal control. The primer information is listed in Table S1.

### Statistical analyses

Statistical analyses were performed using the tests indicated in the corresponding figure legends. Statistical significance was defined as *P* < 0.05. Unless otherwise indicated, at least three independent experiments were performed, > 20 seedlings was harvested per sample.

### Chemical synthesis of HYS analogues

The HYS analogs (HAs) were synthesized in a two-step reaction (Paczal et al., 2006), previously described for HYS synthesis (Zeng et al., 2023). In a 25 mL flask, 2 mmol (1 equiv) of intended acetamide, 2.2 mmol (1.1 equiv) phenol derivate, 2.2 mmol (1.1 equiv) K2CO3 and 0.2 mmol (0.1 equiv) KI were dissolved in 15 mL acetone. The resulting mixture was heated to reflux temperature for 16 hours, after which it was quenched with 5 ml H2O and 5 mL EtOAc, and diluted with 40 mL ethyl acetate. The solution was washed twice with water, and the organic phase was dried over MgSO4 and evaporated. The resulting solid was recrystallized from chloroform, delivering clear crystals. Impurities were removed by filtration over celite. HA13,HA14, HA15 and HA19 contained a sediment (putative hydrolysis product) that could be removed by warm filtration. Impurities in HA21 and HA32 were removed by dissolving in diethylether and 4 washes in 1 M NaOH.

All chemicals used in the rooting experiments were dissolved in dimethylsulfoxide (DMSO) at a stock solution concentration of 10 mM and stored at −20°C.

### Chemical synthesis of acetamides

The different acetamides were generated by coupling chloroacetate chloride to aniline dervatives with corresponding R-groups. In a dry 50 ml flask, placed under inert atmosphere, 15 mmol (1 equiv) aniline derivate and 16.5 mmol (1.1 equiv) triethylamine were dissolved in 20 mL of dry dichloromethane under inert atmosphere. The mixture was cooled to 0 °C and over a period of 5 min, 16.5 mmol (1.1 equiv) chloroacetylchloride was added dropwise. The mixture was stirred for 15 min at 0 °C and subsequently for 3 h at room temperature. Workup was performed by extraction from saturated aqueous NaHCO_3_ (75 mL) by means of dichloromethane (3 x 40 mL). After drying over MgSO_4_ and evaporation of the volatiles under vacuum, crystals were obtained. To remove impurities (possibly triethylammoniumhydrochloride salts), the crystals were dissolved in EtOAc and filtered over a sintered glass filter with a layer of 1 cm celite. Evaporation of the volatiles in the filtrate yielded pure acetamide. For acetamide nr 29, this washing step was repeated. Acetamide 30 was not stable during the reaction and was purified using flash chromatography (PE/EtOAc 7:3; Rf 0.32) to obtain 150 mg pure acetamide (yield = 12 %).

## Supplementary Data

### Spectral data for 2-chloro-N-phenylacetamide (Amide 25)

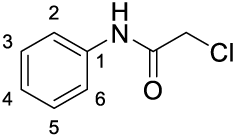

**^1^H-NMR** (400 MHz, CDCl_3_): δ 4.16 (2H, s, CH_2_); 7.16 (1H, t, J=7.4 Hz, C^4^H); 7.34 (2H, t, J=8.0 Hz, C^3^H + C^5^H); 7.53 (2H, d, J=7.6 Hz, C^2^H + C^6^H); 8.29 (1H, br. s, NH). **^13^C-NMR** (100 MHz, CDCl_3_): δ 42.9 (CH_2_); 120.2 (C^2^H + C^6^H); 125.3 (C^4^H); 129.1 (C^3^H + C^5^H); 136.7 (C^1^); 163.9 (NHCO). **IR** (cm^-1^) ν_max_: 1668 (CO). Yield = 88 %.

### Spectral data for 2-chloro-N-(o-tolyl)acetamide 26

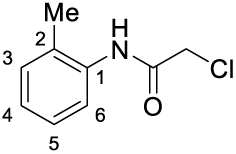

**^1^H-NMR** (400 MHz, CDCl_3_): δ 2.27 (3H, s, C^2^<u>Me</u>); 4.19 (2H, s, CH_2_); 7.10 (1H, t, J=7.4 Hz, C^5^H); 7.15-7.25 (2H, m, C^3^H + C^4^H); 7.83 (1H, d, J=7.9 Hz, C^6^H); 8.23 (1H, br. s, NH). **^13^C-NMR** (100 MHz, CDCl_3_): δ 17.5 (C^2^<u>Me</u>); 43.2 (CH_2_); 122.6 (C^6^H); 125.9 (C^5^H); 126.9 (C^4^H); 129.2 (C^2^); 130.6 (C^3^H); 134.7 (C^1^); 163.9 (NHCO). **IR** (cm^-1^) ν_max_: 1660 (CO); 3265 (NH). Yield = 95 %.

### Spectral data for 2-chloro-N-(4-iodophenyl)acetamide 27

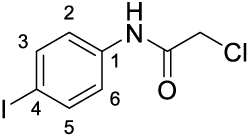

**^1^H-NMR** (400 MHz, CDCl_3_): δ 4.18 (2H, s, CH_2_); 7.34 (2H, d, J=8.8 Hz, C^2^H + C^6^H); 7.67 (2H, d, J=8.8 Hz, C^3^H + C^5^H); 8.21 (1H, br. s, NH). **^13^C-NMR** (100 MHz, CDCl_3_): δ 42.9 (CH_2_); 88.7 (C^4^I); 121.9 (C^2^H + C^6^H); 136.5 (C^1^); 138.1 (C^3^H + C^5^H); 163.8 (NHCO). **IR** (cm^-1^) ν_max_: 1660 (CO); 3261 (NH). Yield = 90 %.

### Spectral data for 2-chloro-N-(4-iodo-2-methylphenyl)acetamide 28

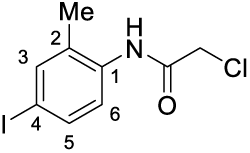

**^1^H-NMR** (400 MHz, CDCl_3_): δ 2.25 (3H, s, C^2^<u>Me</u>); 4.22 (2H, s, CH_2_); 7.54 (1H, d, J=8.0 Hz, C^5^H); 7.55 (1H, s, C^3^H, C^3^H); 7.69 (1H, d, J=8.0 Hz, C^6^H); 8.19 (1H, br. s, NH). **^13^C-NMR** (100 MHz, CDCl_3_): δ 17.1 (C^2^<u>Me</u>); 43.2 (CH_2_); 89.7 (C^4^I); 123.8 (C^6^H); 131.0 (C^2^); 134.6 (C^1^); 136.0 (C^5^H); 139.2 (C^3^H); 163.7 (NHCO). **IR** (cm^-1^) ν_max_: 1662 (CO); 3248 (NH). Yield = 91 %.

### Spectral data for Methyl 2-(2-chloroacetamido)-5-iodobenzoate 29

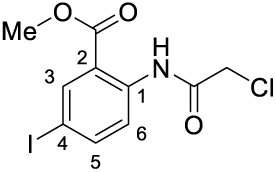

**^1^H-NMR** (400 MHz, CDCl_3_): δ 3.96 (3H, s, O<u>Me</u>); 4.20 (2H, s, CH_2_); 7.83 (1H, d, J=8.9 Hz, C^5^H); 8.36 (1H, s, C^3^H); 8.49 (1H, d, J=8.9 Hz, C^6^H); 11.80 (1H, br. s, NH). **^13^C-NMR** (100 MHz, CDCl_3_): δ 43.2 (CH_2_); 52.8 (O<u>Me</u>); 86.3 (C^4^I); 117.6 (C^2^); 122.2 (C^6^H); 139.5 (C^3^H); 139.9 (C^1^); 143.1 (C^5^H); 165.3 (NHCO). 167.0 (C^2^<u>C</u>O). **MS** (ESI): m/z (%): 354 (M+H^+^, 100); 376 (M+Na^+^, 45). **IR** (cm^-1^) ν_max_: 1672 (CO); 1701 (CO); 3226 (NH). Yield = 76 %.

### Spectral data for 2-chloro-N-(4-iodo-2-nitrophenyl)acetamide 30

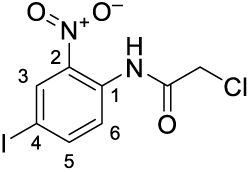

**^1^H-NMR** (400 MHz, CDCl_3_): δ 4.24 (2H, s, CH_2_); 7.95 (1H, dd, J=8.9, 2.1 Hz, C^5^H); 8.56 (1H, d, J=8.9 Hz, C^6^H); 8.56 (1H, d, J=2.1 Hz, C^3^H); 11.31 (1H, br. s, NH). **^13^C-NMR** (100 MHz, CDCl_3_): δ 43.1 (CH_2_); 86.1 (C^4^I); 123.5 (C^6^H); 133.3 (C^1^); 134.2 (C^3^H); 137.1 (C^2^); 144.5 (C^5^H); 165.3 (NHCO). **MS** (ESI): m/z (%): 105 (40); 225 (100); 290 (25); 317 (25); 362 (M+Na^+^, 10); 393 (30). **IR** (cm^-1^) ν_max_: 1681 (CO); 3288 (NH). Rf = 0.32 (Pe/EtOAc 7/3). Yield = 12 %.

### Spectral data for Methyl 4-(2-chloroacetamido)-3-methylbenzoate 31

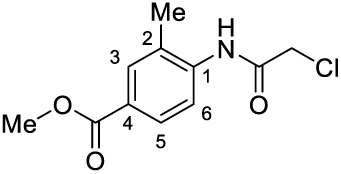

**^1^H-NMR** (400 MHz, CDCl_3_): δ 2.35 (3H, s, C^2^<u>Me</u>); 3.90 (3H, s, O<u>Me</u>); 4.25 (2H, s, CH_2_); 7.90 (1H, s, C^3^H); 7.91 (1H, d, J=8.3 Hz, C^5^H); 8.16 (1H, d, J=8.3 Hz, C^6^H); 8.17 (1H, br. s, NH). **^13^C-NMR** (100 MHz, CDCl_3_): δ 17.4 (C^2^<u>Me</u>); 43.2 (CH_2_); 52.1 (O<u>Me</u>); 120.5 (C^6^H); 126.5 (C^1^); 127.4 (C^2^); 128.7 (C^5^H); 131.9 (C^3^H); 139.0 (C^4^); 163.7 (NHCO). 166.6 (C^4^<u>C</u>O). **MS** (ESI): m/z (%): 242 (M+H^+^, 100); 264 (M+Na^+^, 25). **IR** (cm^-1^) ν_max_: 1666 (CO); 1720 (CO); 2941; 3265 (NH). Yield = 56 %.

### Spectral data for 2-chloro-N-(4-methoxy-2-methylphenyl)acetamide 32

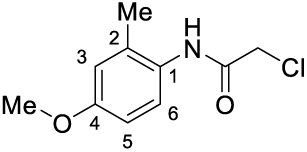

**^1^H-NMR** (400 MHz, CDCl_3_): δ 2.25 (3H, s, C^2^<u>Me</u>); 3.78 (3H, s, O<u>Me</u>); 4.20 (2H, s, CH_2_); 6.72-6.77 (2H, m, C^3^H + C^5^H); 7.55-7.59 (1H, m, C^6^H); 8.07 (1H, br. s, NH). **^13^C-NMR** (100 MHz, CDCl_3_): δ 17.9 (C^2^<u>Me</u>); 43.0 (CH_2_); 55.4 (O<u>Me</u>); 111.7 (C^5^H); 116.1 (C^3^H); 125.0 (C^6^H); 127.5 (C^1^); 132.4 (C^2^); 157.6 (C^4^O); 164.1 (NHCO). **MS** (ESI): m/z (%): 214 (M+H^+^, 100). **IR** (cm^-1^) ν_max_: 1664 (CO); 2833; 3005; 3267 (NH). Yield = 57 %.

### Spectral data for 2-chloro-N-(4-fluoro-2-methylphenyl)acetamide 33

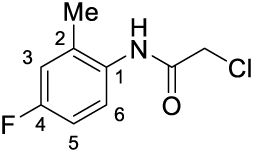

**^1^H-NMR** (400 MHz, CDCl_3_): δ 2.28 (3H, s, C^2^<u>Me</u>); 4.23 (2H, s, CH_2_); 6.88-6.96 (2H, m, C^3^H + C^5^H); 7.70-7.77 (1H, m, C^6^H); 8.12 (1H, br. s, NH). **^13^C-NMR** (100 MHz, CDCl_3_): δ 17.7 (C^2^<u>Me</u>); 43.0 (CH_2_); 113.5 (1C, d, J=22.4 Hz, C^3^H); 117.2 (1C, d, J=22.4 Hz, C^5^H); 124.8 (1C, d, J=8.2 Hz, C^6^H); 130.5 (1C, d, J=2.8 Hz, C^1^); 132.3 (1C, d, J=8.2 Hz, C^2^); 160.3 (1C, d, J=245.1 Hz, C^4^F); 164.0 (NHCO). **MS** (ESI): m/z (%): 202 (M+H^+^, 100); 224 (M+Na^+^, 15). **IR** (cm^-1^) ν_max_: 1662 (CO); 3253 (NH). Yield = 76 %.

### Spectral data for 2-phenoxy-N-phenylacetamide HA1

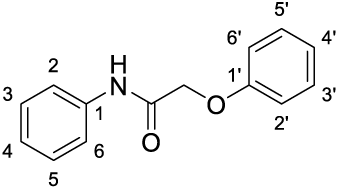

**^1^H-NMR** (400 MHz, CDCl_3_): δ 4.49 (2H, s, OCH_2_); 6.92 (2H, d, J=7.8 Hz, C^2’^H + C^6’^H); 7.01 (1H, t, J=7.4 Hz, C^4’^H); 7.10 (1H, t, J=7.4 Hz, C^4^H); 7.29 (4H, t, J=7.2 Hz, C^3^H + C^3’^H + C^5^H + C^5’^H); 7.57 (2H, d, J=7.8 Hz, C^2^H + C^6^H); 8.38 (1H, br. s, NH). **^13^C-NMR** (100 MHz, CDCl_3_): δ 67.6 (OCH_2_); 114.9 (C^2’^H+C^6’^H); 120.3 (C^2^H+C^6^H); 122.4 (C^4’^H); 124.9 (C^4^H); 129.1 (C^3^H+C^5^H); 129.9 (C^3’^H+C^5’^H); 137.0 (C^1^); 157.1 (C^1’^O); 166.4 (NHCO). **MS** (ESI): m/z (%): 228 (M+H^+^, 100); 250 (M+Na^+^, 40). **IR** (cm^-1^) ν_max_: 1661 (CO). Yield = 75 %.

### Spectral data for N-phenyl-2-(o-tolyloxy)acetamide HA2

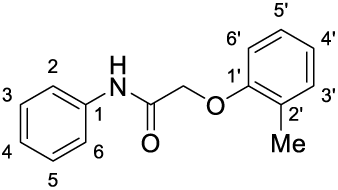

**^1^H-NMR** (400 MHz, CDCl_3_): δ 2.35 (3H, s, C^2’^<u>Me</u>); 4.55 (2H, s, OCH_2_); 6.80 (1H, d, J=8.0 Hz, C^6’^H); 6.95 (1H, t, J=7.4 Hz, C^4’^H); 7.10-7.22 (3H, m, C^3’^H + C^4^H + C^5’^H); 7.33 (2H, t, J=7.9 Hz, C^3^H + C^5^H); 7.57 (2H, d, J=8.2 Hz, C^2^H + C^6^H); 8.34 (1H, br. s, NH). **^13^C-NMR** (100 MHz, CDCl_3_): δ 16.4 (C^2’^<u>Me</u>); 67.8 (OCH_2_); 111.9 (C^6’^H); 120.0 (C^2^H + C^6^H); 122.2 (C^4’^H); 124.9 (C^4^H); 126.5 (C^2’^); 127.4 (C^5’^H); 129.2 (C^3^H + C^5^H); 131.2 (C^3’^H); 137.0 (C^1^); 155.3 (C^1’^O); 166.4 (NHCO). **MS** (ESI): m/z (%): 242 (M+H^+^, 100); 264 (M+Na^+^, 40). **IR** (cm^-1^) ν_max_: 1676 (CO); 3356 (NH). Yield = 75 %.

### Spectral data for 2-(4-chlorophenoxy)-N-phenylacetamide HA3

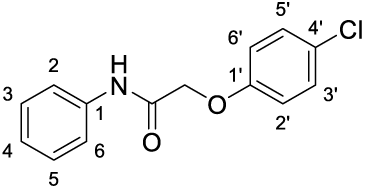

**^1^H-NMR** (400 MHz, CDCl_3_): δ 4.56 (2H, s, OCH_2_); 6.92 (2H, d, J=9.0 Hz, C^2’^H + C^6’^H); 7.15 (1H, t, J=7.4 Hz, C^4^H); 7.29 (2H, d, J=9.0 Hz, C^3’^H + C^5’^H); 7.35 (2H, t, J=8.0 Hz, C^3^H + C^5^H); 7.57 (2H, d, J=7.6 Hz, C^2^H + C^6^H); 8.22 (1H, br. s, NH). **^13^C-NMR** (100 MHz, CDCl_3_): δ 67.9 (OCH_2_); 116.2 (C^2’^H + C^6’^H); 120.2 (C^2^H + C^6^H); 125.0 (C^4^H); 127.5 (C^4’^Cl); 129.1 (C^3^H + C^5^H); 129.8 (C^3’^H + C^5’^H); 136.7 (C^1^); 155.6 (C^1’^O); 165.8 (NHCO). **MS** (ESI): m/z (%): 262 (M+H^+^, 100); 264 (M+H^+^, 30); 284 (M+Na^+^, 55); 286 (M+Na^+^, 15). **IR** (cm^-1^) ν_max_: 1674 (CO); 3356 (NH). Yield = 96 %.

### Spectral data for 2-(4-chloro-2-methylphenoxy)-N-phenylacetamide HA4

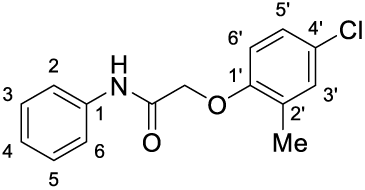

**^1^H-NMR** (400 MHz, CDCl_3_): δ 2.33 (3H, s, C^2’^<u>Me</u>); 4.56 (2H, s, OCH_2_); 6.75 (1H, d, J=8.6 Hz, C^6’^H); 7.11-7.20 (3H, m, C^3’^H + C^4^H + C^5’^H); 7.35 (2H, t, J=7.9 Hz, C^3^H + C^5^H); 7.57 (2H, d, J=7.6 Hz, C^2^H + C^6^H); 8.25 (1H, br. s, NH). **^13^C-NMR** (100 MHz, CDCl_3_): δ 15.8 (C^2’^<u>Me</u>); 67.9 (OCH_2_); 113.1 (C^6’^H); 120.0 (C^2^H + C^6^H); 125.0 (C^5’^H); 127.0 (C^3’^H); 127.1 (C^2’^); 128.4 (C^4’^Cl); 129.2 (C^3^H + C^5^H); 131.0 (C^4^H); 136.8 (C^1^); 153.9 (C^1’^O); 165.9 (NHCO). **MS** (ESI): m/z (%): 102 (30); 276 (M+H^+^, 100); 278 (M+H^+^, 30); 298 (M+Na^+^, 50); 300 (M+Na^+^, 10); 304 (30);. **IR** (cm^-1^) ν_max_: 1672 (CO); 3356 (NH). Yield = 89 %.

### Spectral data for 2-phenoxy-N-(o-tolyl)acetamide HA5

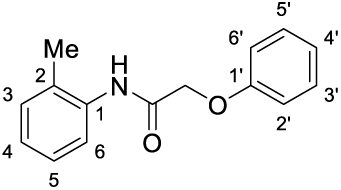

**^1^H-NMR** (400 MHz, CDCl_3_): δ 2.21 (3H, s, C^2^<u>Me</u>); 4.64 (2H, s, OCH_2_); 6.98 (2H, d, J=7.6 Hz, C^2’^H + C^6’^H); 7.05 (1H, t, J=7.4 Hz, C^4’^H); 7.09 (1H, t, J=6.8 Hz, C^5^H); 7.18 (1H, d, J=7.9 Hz, C^3^H); 7.22 (1H, t, J=7.8 Hz, C^4^H); 7.34 (2H, dd, J=7.6, 7.4 Hz, C^3’^H + C^5’^H); 7.96 (1H, d, J=8.0 Hz, C^6^H); 8.25 (1H, br. s, NH). **^13^C-NMR** (100 MHz, CDCl_3_): δ 17.4 (C^2^<u>Me</u>); 67.6 (OCH_2_); 114.7 (C^2’^H+C^6’^H); 122.4 (C^6^H); 122.5 (C^4’^H); 125.4 (C^5^H); 127.0 (C^4^H); 128.7 (C^2^); 130.0 (C^3’^H+C^5’^H); 130.6 (C^3^H); 134.8 (C^1^); 157.0 (C^1’^O); 166.3 (NHCO). **MS** (ESI): m/z (%): 242 (M+H^+^, 100); 264 (M+Na^+^, 15). **IR** (cm^-1^) ν_max_: 1667 (CO). Yield = 60 %.

### Spectral data for N-(o-tolyl)-2-(o-tolyloxy)acetamide HA6

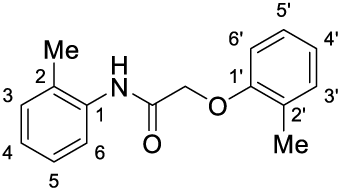

**^1^H-NMR** (400 MHz, CDCl_3_): δ 2.26 (3H, s, C^2^<u>Me</u>); 2.36 (3H, s, C^2’^<u>Me</u>); 4.64 (2H, s, OCH_2_); 6.85 (1H, d, J=8.4 Hz, C^6’^H); 6.97 (1H, t, J=7.4 Hz, C^4’^H); 7.09 (1H, td, J=7.5 Hz, C^5^H); 7.19 (1H, d, J=7.0 Hz, C^3^H); 7.20 (1H, t, J=7.2 Hz, C^5’^H); 7.21 (1H, d, J=7.5 Hz, C^3’^H); 7.24 (1H, t, J=8.5 Hz, C^4^H); 8.08 (1H, d, J=8.1 Hz, C^6^H); 8.32 (1H, br. s, NH). **^13^C-NMR** (100 MHz, CDCl_3_): δ 16.5 (C^2’^<u>Me</u>); 17.5 (C^2^<u>Me</u>); 67.5 (OCH_2_); 111.4 (C^6’^H); 121.9 (C^6^H); 122.1 (C^4’^H); 125.1 (C^5^H); 126.3 (C^2’^); 127.0 (C^4^H); 127.4 (C^5’^H); 127.8 (C^2^); 130.5 (C^3^H); 131.2 (C^3’^H); 135.0 (C^1^); 155.1 (C^1’^O); 166.2 (NHCO). **MS** (ESI): m/z (%): 256 (M+H^+^, 100); 278 (M+Na^+^, 15). **IR** (cm^-1^) ν_max_: 1682 (CO); 3410 (NH). Yield = 67 %.

### Spectral data for 2-(4-chlorophenoxy)-N-(o-tolyl)acetamide HA7

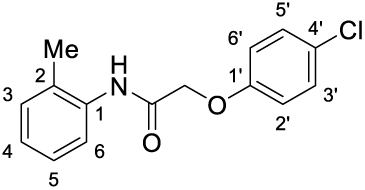

**^1^H-NMR** (400 MHz, CDCl_3_): δ 2.21 (3H, s, C^2^<u>Me</u>); 4.59 (2H, s, OCH_2_); 6.90 (2H, d, J=8.9 Hz, C^2’^H + C^6’^H); 7.09 (1H, t, J=7.4 Hz, C^5^H); 7.16-7.25 (2H, m, C^3^H + C^4^H); 7.29 (2H, d, J=8.9 Hz, C^3’^H + C^5’^H); 7.93 (1H, d, J=8.0 Hz, C^6^H); 8.18 (1H, br. s, NH). **^13^C-NMR** (100 MHz, CDCl_3_): δ 17.5 (C^2^<u>Me</u>); 67.9 (OCH_2_); 116.0 (C^2’^H+ C^6’^H); 122.5 (C^6^H); 125.5 (C^5^H); 127.0 (C^4^H); 127.5 (C^4’^Cl); 128.7 (C^2^); 129.9 (C^3’^H+ C^5’^H); 130.6 (C^3^H); 134.7 (C^1^); 155.6 (C^1’^O); 165.7 (NHCO). **MS** (ESI): m/z (%): 276 (M+H^+^, 100); 278 (M+H^+^, 35); 298 (M+Na^+^, 40); 300 (M+Na^+^, 10). **IR** (cm^-1^) ν_max_: 1670 (CO); 3262 (NH). Yield = 79 %.

### Spectral data for 2-(4-chloro-2-methylphenoxy)-N-(o-tolyl)acetamide HA8

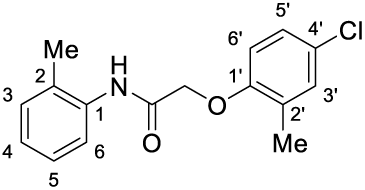

**^1^H-NMR** (400 MHz, CDCl_3_): δ 2.26 (3H, s, C^2^<u>Me</u>); 2.32 (3H, s, C^2’^<u>Me</u>); 4.60 (2H, s, OCH_2_); 6.76 (1H, d, J=8.6 Hz, C^6’^H); 7.08 (1H, td, J=7.5, 0.9 Hz, C^5^H); 7.15 (1H, d, J=8.6, C^5’^H); 7.17-7.21 (2H, m, C^3’^H + C^3^H); 7.24 (1H, t, J=7.8 Hz, C^4^H); 8.06 (1H, d, J=8.1 Hz, C^6^H); 8.24 (1H, br. s, NH). **^13^C-NMR** (100 MHz, CDCl_3_): δ 16.4 (C^2’^<u>Me</u>); 17.5 (C^2^<u>Me</u>); 67.8 (OCH_2_); 112.5 (C^6’^H); 121.9 (C^6^H); 125.3 (C^5^H); 126.9 (C^2’^); 127.0 (C^5’^H); 127.1 (C^4^H); 127.8 (C^2^); 128.2 (C^4’^Cl); 130.6 (C^3^H); 131.0 (C^3’^H); 134.9 (C^1^); 153.7 (C^1’^O); 165.7 (NHCO). **MS** (ESI): m/z (%): 290 (M+H^+^, 100); 292 (M+H^+^, 35); 312 (M+Na^+^, 30); 314 (M+Na^+^, 15). **IR** (cm^-1^) ν_max_: 1695 (CO); 3416 (NH). Yield = 70 %.

### Spectral data for N-(4-iodophenyl)-2-phenoxyacetamide HA9

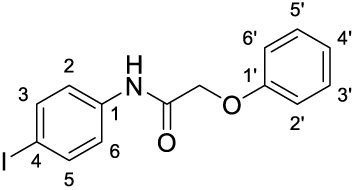

**^1^H-NMR** (400 MHz, CDCl_3_): δ 4.59 (2H, s, OCH_2_); 6.98 (2H, d, J=7.8 Hz, C^2’^H + C^6’^H); 7.07 (1H, t, J=7.4 Hz, C^4’^H); 7.35 (2H, t, J=7.9 Hz, C^3’^H + C^5’^H); 7.38 (2H, d, J=8.8 Hz, C^2^H + C^6^H); 7.65 (2H, d, J=8.8 Hz, C^3^H + C^5^H); 8.27 (1H, br. s, NH). **^13^C-NMR** (100 MHz, CDCl_3_): δ 67.6 (OCH_2_); 88.2 (C^4^I); 114.8 (C^2’^H+C^6’^H); 121.9 (C^2^H+C^6^H); 122.6 (C^4’^H); 130.0 (C^3’^H+C^5’^H); 136.6 (C^1^); 138.0 (C^3^H+C^5^H); 156.9 (C^1’^O); 166.3 (NHCO). **MS** (ESI): m/z (%): 102 (30); 304 (40); 354 (M+H^+^, 100); 376 (M+Na^+^, 55). **IR** (cm^-1^) ν_max_: 1670 (CO); 3360 (NH). Yield = 98 %.

### Spectral data for N-(4-iodophenyl)-2-(o-tolyloxy)acetamide HA10

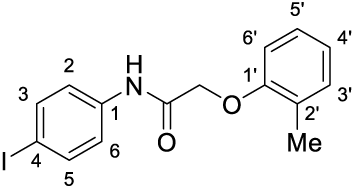

**^1^H-NMR** (400 MHz, CDCl_3_): δ 2.35 (3H, s, C^2’^<u>Me</u>); 4.57 (2H, s, OCH_2_); 6.81 (1H, d, J=8.0 Hz, C^6’^H); 6.97 (1H, t, J=7.4 Hz, C^4’^H); 7.16-7.23 (2H, m, C^3’^H + C^5’^H); 7.36 (2H, d, J=8.7 Hz, C^2^H + C^6^H); 7.64 (2H, d, J=8.7 Hz, C^3^H + C^5^H); 8.33 (1H, br. s, NH). **^13^C-NMR** (100 MHz, CDCl_3_): δ 16.5 (C^2’^<u>Me</u>); 67.8 (OCH_2_); 88.2 (C^4^I); 112.0 (C^6’^H); 121.7 (C^2^H+C^6^H); 122.4 (C^4’^H); 126.5 (C^2’^); 127.4 (C^5’^H); 131.3 (C^3’^H); 136.7 (C^1^); 138.1 (C^3^H+C^5^H); 155.2 (C^1’^O); 166.5 (NHCO). **MS** (ESI): m/z (%): 102 (30); 368 (M+H^+^, 100); 390 (M+Na^+^, 35). **IR** (cm^-1^) ν_max_: 1688 (CO); 3389 (NH). Yield = 92 %.

### Spectral data for 2-(4-chlorophenoxy)-N-(4-iodophenyl)acetamide HA11

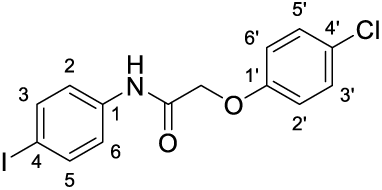

**^1^H-NMR** (400 MHz, CDCl_3_): δ 4.56 (2H, s, OCH_2_); 6.91 (2H, d, J=9.0 Hz, C^2’^H + C^6’^H); 7.30 (2H, d, J=9.0 Hz, C^3’^H + C^5’^H); 7.37 (2H, d, J=8.7 Hz, C^2^H + C^6^H); 7.65 (2H, d, J=8.7 Hz, C^3^H + C^5^H); 8.20 (1H, br. s, NH). **^13^C-NMR** (100 MHz, CDCl_3_): δ 67.9 (OCH_2_); 88.4 (C^4^I); 116.2 (C^2’^H+C^6’^H); 121.9 (C^2^+C^6^); 127.7 (C^4’^Cl); 129.9 (C^3’^H+C^5’^H); 136.5 (C^1^); 138.1 (C^3^H+C^5^H); 155.4 (C^1’^O); 165.8 (NHCO). **MS** (ESI): m/z (%): 102 (100); 304 (45); 388 (M+H^+^, 30); 390 (M+H^+^, 10); 410 (M+Na^+^, 35); 412 (M+Na^+^, 10). **IR** (cm^-1^) ν_max_: 1697 (CO); 3372 (NH). Yield = 88 %.

### Spectral data for 2-(4-chloro-2-methylphenoxy)-N-(4-iodophenyl)acetamide HA12

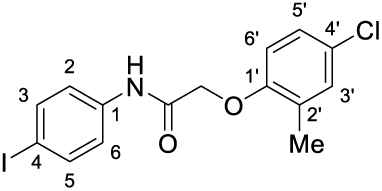

**^1^H-NMR** (400 MHz, CDCl_3_): δ 2.24 (3H, s, C^2’^<u>Me</u>); 4.45 (2H, s, OCH_2_); 6.64 (1H, d, J=8.6 Hz, C^6’^H); 7.05 (1H, dd, J=8.6, 2.0 Hz, C^5’^H); 7.09 (1H, d, J=2.0 Hz, C^3’^H); 7.27 (2H, d, J=8.6 Hz, C^2^H + C^6^H); 7.55 (2H, d, J=8.6 Hz, C^3^H + C^5^H); 8.16 (1H, br. s, NH). **^13^C-NMR** (100 MHz, CDCl_3_): δ 15.3 (C^2’^<u>Me</u>); 67.0 (OCH_2_); 87.2 (C^4^I); 112.0 (C^6’^H); 120.7 (C^2^+C^6^); 126.0 (C^5’^H); 126.1 (C^4’^Cl); 127.4 (C^2’^); 130.0 (C^3’^H); 135.5 (C^1^); 137.0 (C^3^H+C^5^H); 152.7 (C^1’^O); 164.9 (NHCO). **MS** (ESI): m/z (%): 102 (55); 304 (100); 402 (M+H^+^, 30); 404 (M+H^+^, 10); 424 (M+Na^+^, 15); 424 (M+Na^+^, 5). **IR** (cm^-1^) ν_max_: 1670 (CO); 3302 (NH). Yield = 96 %.

### Spectral data for N-(4-iodo-2-methylphenyl)-2-phenoxyacetamide HA13

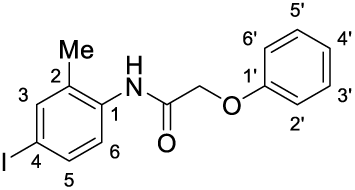

**^1^H-NMR** (400 MHz, CDCl_3_): δ 2.18 (3H, s, C^2^<u>Me</u>); 4.65 (2H, s, OCH_2_); 6.98 (2H, d, J=7.8 Hz, C^2’^H + C^6’^H); 7.07 (1H, t, J=7.4 Hz, C^4’^H); 7.36 (2H, dd, J=7.8, 7.4 Hz, C^3’^H + C^5’^H); 7.53 (1H, s, C^3^H); 7.54 (1H, d, J=9.0 Hz, C^5^H); 7.81 (1H, d, J=9.0 Hz, C^6^H); 8.22 (1H, br. s, NH). **^13^C-NMR** (100 MHz, CDCl_3_): δ 17.0 (C^2^<u>Me</u>); 67.6 (OCH_2_); 89.1 (C^4^I); 114.7 (C^2’^H + C^6’^H); 122.6 (C^4’^H); 123.7 (C^6^H); 130.0 (C^3’^H + C^5’^H); 130.5 (C^2^); 134.8 (C^1^); 135.9 (C^5^H); 139.1 (C^3^H); 156.8 (C^1’^O); 166.2 (NHCO). **MS** (ESI): m/z (%): 368 (M+H^+^, 100); 390 (M+Na^+^, 25). **IR** (cm^-1^) ν_max_: 1693 (CO); 3402 (NH). Yield = 46 %.

### Spectral data for N-(4-iodo-2-methylphenyl)-2-(o-tolyloxy)acetamide HA14

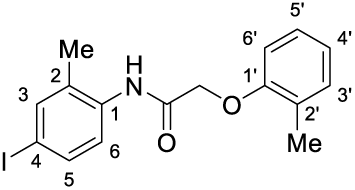

**^1^H-NMR** (400 MHz, CDCl_3_): δ 2.22 (3H, s, C^2^<u>Me</u>); 2.35 (3H, s, C^2’^<u>Me</u>); 4.64 (2H, s, OCH_2_); 6.84 (1H, d, J=8.4 Hz, C^6’^H); 6.98 (1H, t, J=7.2 Hz, C^4’^H); 7.18-7.24 (2H, m, C^3’^H + C^5’^H); 7.54 (1H, s, C^3^H); 7.55 (1H, d, J=8.2 Hz, C^5^H); 7.91 (1H, d, J=8.2 Hz, C^6^H); 8.30 (1H, br. s, NH). **^13^C-NMR** (100 MHz, CDCl_3_): δ 16.5 (C^2’^<u>Me</u>); 17.1 (C^2^<u>Me</u>); 67.5 (OCH_2_); 88.8 (C^4^I); 111.4 (C^6’^H); 122.2 (C^4’^H); 123.3 (C^6^H); 126.2 (C^2’^); 127.4 (C^5’^H); 129.8 (C^2^); 131.3 (C^3’^H); 134.9 (C^1^); 136.0 (C^5^H); 139.1 (C^3^H); 154.9 (C^1’^O); 166.2 (NHCO). **MS** (ESI): m/z (%): 382 (M+H^+^, 100); 404 (M+Na^+^, 35). **IR** (cm^-1^) ν_max_: 1697 (CO); 3406 (NH). Yield = 39 %.

### Spectral data for 2-(4-chlorophenoxy)-N-(4-iodo-2-methylphenyl)acetamide HA15

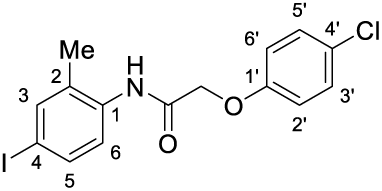

**^1^H-NMR** (400 MHz, CDCl_3_): δ 2.19 (3H, s, C^2^<u>Me</u>); 4.62 (2H, s, OCH_2_); 6.92 (2H, d, J=9.0 Hz, C^3’^H + C^5’^H); 7.31 (2H, d, J=9.0 Hz, C^2’^H + C^6’^H); 7.54 (1H, s, C^3^H); 7.55 (1H, d, J=9.1 Hz, C^5^H); 7.78 (1H, d, J=9.1 Hz, C^6^H); 8.14 (1H, br. s, NH). **^13^C-NMR** (100 MHz, CDCl_3_): δ 17.0 (C^2^<u>Me</u>); 67.9 (OCH_2_); 89.3 (C^4^I); 116.0 (C^2’^H + C^6’^H); 123.8 (C^6^H); 127.7 (C^4’^Cl); 129.9 (C^3’^H + C^5’^H); 130.5 (C^2^); 134.6 (C^1^); 136.0 (C^5^H); 139.2 (C^3^H); 155.4 (C^1’^O); 165.7 (NHCO). **MS** (ESI): m/z (%): 102 (65); 177 (35); 304 (25); 402 (M+H^+^, 100); 404 (M+H^+^, 35); 424 (M+Na^+^, 65); 426 (M+Na^+^, 25). **IR** (cm^-1^) ν_max_: 1701 (CO); 3406 (NH). Yield = 69 %.

### Spectral data for 2-(4-fluoro-2-methylphenoxy)-N-(4-iodo-2-methylphenyl)acetamide HA17

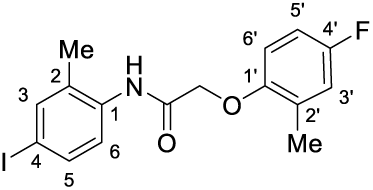

**^1^H-NMR** (400 MHz, CDCl_3_): δ 2.22 (3H, s, C^2^<u>Me</u>); 2.33 (3H, s, C^2’^<u>Me</u>); 4.59 (2H, s, OCH_2_); 6.74-6.80 (1H, m, C^6’^H); 6.84-6.91 (1H, m, C^5’^H); 6.91-6.97 (1H, m, C^3’^H); 7.54 (1H, s, C^3^H); 7.55 (1H, d, J=9.1 Hz, C^6^H); 7.89 (1H, d, J=9.1 Hz, C^5^H); 8.25 (1H, br. s, NH). **^13^C-NMR** (100 MHz, CDCl_3_): δ 16.6 (C^2’^<u>Me</u>); 17.1 (C^2^<u>Me</u>); 68.2 (OCH_2_); 88.9 (C^4^I); 112.5 (d, J^CF^=8.7 Hz, C^6’^H); 113.1 (d, J^CF^=23.1 Hz, C^5’^H); 118.1 (d, J^CF^=23.1 Hz, C^3’^H); 123.4 (C^5^H); 128.2 (d, J^CF^=7.9 Hz, C^2’^); 129.8 (C^2^); 134.9 (C^1^); 136.0 (C^6^H); 139.1 (C^3^H); 151.1 (d, J^CF^=2.1 Hz, C^1’^O); 157.7 (d, J^CF^=240.4 Hz, C^4’^F); 165.9 (NHCO). **MS** (ESI): m/z (%): 400 (M+H^+^, 100); 422 (M+Na^+^, 25). **HRMS** (ESI): Calculated for C_16_H_15_FINO_2_^+^: 400.0204265 [M + H]^+^, found: 400.02192. **IR** (cm^-1^) ν_max_: 1662 (CO); 3410 (NH). Yield = 56 %.

### Spectral data for 2-(4-bromo-2-methylphenoxy)-N-(4-iodo-2-methylphenyl)acetamide HA18

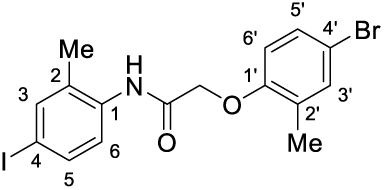

**^1^H-NMR** (400 MHz, CDCl_3_): δ 2.21 (3H, s, C^2^<u>Me</u>); 2.31 (3H, s, C^2’^<u>Me</u>); 4.58 (2H, s, OCH_2_); 6.70 (1H, d, J=8.6 Hz, C^6’^H); 7.30 (1H, d, J=8.6 Hz, C^5’^H); 7.32 (1H, s, C^3’^H); 7.52 (1H, d, J=9.1 Hz, C^6^H); 7.87 (1H, d, J=9.1 Hz, C^5^H); 8.21 (1H, br. s, NH). **^13^C-NMR** (100 MHz, CDCl_3_): δ 16.4 (C^2’^<u>Me</u>); 17.1 (C^2^<u>Me</u>); 67.7 (OCH_2_); 89.0 (C^4^I); 113.0 (C^6’^H); 114.5 (C^2’^); 123.3 (C^6^H); 129.8 (C^2^); 130.0 (C^5’^H); 133.9 (C^3’^H); 134.8 (C^1^); 136.0 (C^5^H); 138.2 (C^4’^Br); 139.1 (C^3^H); 154.1 (C^1’^O); 165.6 (NHCO). **MS** (ESI): m/z (%): 461 (M+H^+^, 100); 483 (M+Na^+^, 55). **HRMS** (ESI): Calculated for C_16_H_15_BrINO_2_^+^: 459.9403609 [M + H]^+^, found: 459.93969. **IR** (cm^-1^) ν_max_: 1662 (CO). Yield = 60 %.

### Spectral data for 2-(4-acetyl-2-methylphenoxy)-N-(4-iodo-2-methylphenyl)acetamide HA19

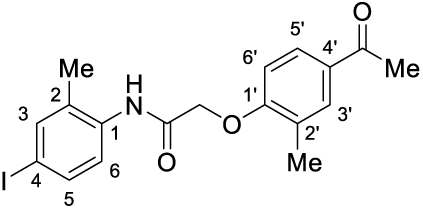

**^1^H-NMR** (400 MHz, CDCl_3_): δ 2.22 (3H, s, C^2^<u>Me</u>); 2.40 (3H, s, C^2’^<u>Me</u>); 2.57 (3H, s, CO<u>Me</u>); 4.69 (2H, s, OCH_2_); 6.88 (1H, d, J=8.9 Hz, C^6’^H); 7.50-7.57 (2H, m, C^3^H + C^6^H); 7.84 (1H, d, J=8.9 Hz, C^5’^H); 7.85 (1H, s, C^3’^H); 7.89 (1H, d, J=9.0 Hz, C^5^H); 8.21 (1H, br. s, NH). **^13^C-NMR** (100 MHz, CDCl_3_): δ 16.6 (C^2’^<u>Me</u>); 17.1 (C^2^<u>Me</u>); 26.4 (CO<u>Me);</u> 67.4 (OCH_2_); 89.0 (C^4^I); 110.6 (C^6’^H); 123.3 (C^5^H); 126.4 (C^2’^); 128.8 (C^5’^H); 129.8 (C^2^); 131.4 (C^3’^H); 131.5 (C^4’^); 134.8 (C^1^); 136.0 (C^6^H); 139.1 (C^3^H); 158.6 (C^1’^O); 165.3 (NHCO). 196.8 (<u>C</u>OMe). **MS** (ESI): m/z (%): 424 (M+H^+^, 100); 446 (M+Na^+^, 70). **HRMS** (ESI): Calculated for C_18_H_18_INO_3_^+^: 424.040413 [M + H]^+^, found: 424.04064. **IR** (cm^-1^) ν_max_: 1662 (CO); 1697 (CO); 3421 (NH). Yield = 89 %.

### Spectral data for N-(4-iodo-2-methylphenyl)-2-(2-methyl-4-(trifluoromethoxy) phenoxy)acetamide HA20

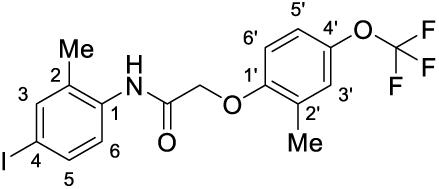

**^1^H-NMR** (400 MHz, CDCl_3_): δ 2.22 (3H, s, C^2^<u>Me</u>); 2.36 (3H, s, C^2’^<u>Me</u>); 4.63 (2H, s, OCH_2_); 6.82 (1H, d, J=8.7 Hz, C^6’^H); 7.08 (1H, d, J=8.7 Hz, C^5’^H); 7.09 (1H, s, C^3’^H); 7.54 (1H, s, C^3^H); 7.55 (1H, d, J=9.2 Hz, C^6^H); 7.89 (1H, d, J=9.2 Hz, C^5^H); 8.21 (1H, br. s, NH). **^13^C-NMR** (100 MHz, CDCl_3_): δ 16.6 (C^2’^<u>Me</u>); 17.1 (C^2^<u>Me</u>); 67.9 (OCH_2_); 89.0 (C^4^I); 112.0 (C^6’^H); 119.9 (C^5’^H); 120.5 (q, J^CF^=256.5 Hz, CF_3_); 123.4 (C^5^H); 124.2 (C^3’^H); 128.1 (C^2’^); 129.8 (C^2^); 134.8 (C^1^); 136.1 (C^6^H); 139.1 (C^3^H); 143.6 (C^4’^Cl); 153.4 (C^1’^O); 165.6 (NHCO). **^19^F-NMR** (376 MHz, CDCl_3_): δ −58.3 (1F, s, C^4’^F) **MS** (ESI): m/z (%): 466 (M+H^+^, 100); 488 (M+Na^+^, 30). **HRMS** (ESI): Calculated for C_17_H_15_F_3_INO_3_^+^: 466.0121475 [M + H]^+^, found: 466.01121. **IR** (cm^-1^) ν_max_: 1697 (CO); 3408 (NH). Yield = 42 %.

### Spectral data for N-(4-iodo-2-methylphenyl)-2-(4-methoxy-2-methylphenoxy)acetamide HA21

**^1^H-NMR** (400 MHz, CDCl_3_): δ 2.22 (3H, s, C^2^<u>Me</u>); 2.32 (3H, s, C^2’^<u>Me</u>); 3.77 (3H, s, O<u>Me</u>); 4.58 (2H, s, OCH_2_); 6.71 (1H, d, J=8.7 Hz, C^6’^H); 6.77 (1H, d, J=8.7 Hz, C^5’^H); 6.78 (1H, s, C^3’^H); 7.54 (1H, s, C^3^H); 7.55 (1H, d, J=8.2 Hz, C^6^H); 7.91 (1H, d, J=8.2 Hz, C^5^H); 8.31 (1H, br. s, NH). **^13^C-NMR** (100 MHz, CDCl_3_): δ 16.7 (C^2’^<u>Me</u>); 17.1 (C^2^<u>Me</u>); 55.6 (O<u>Me</u>); 68.3 (OCH_2_); 88.7 (C^4^I); 111.2 (C^6’^H); 112.7 (C^5’^H); 117.5 (C^3’^H); 123.3 (C^5^H); 127.7 (C^2’^); 129.8 (C^2^); 135.0 (C^1^); 136.0 (C^6^H); 139.1 (C^3^H); 149.2 (C^1’^O); 154.7 (C^4’^); 166.4 (NHCO). **MS** (ESI): m/z (%): 412 (M+H^+^, 100); 434 (M+Na^+^, 20). **HRMS** (ESI): Calculated for C_17_H_18_INO_3_ ^+^: 412.040413 [M + H]^+^, found: 412.0395. **IR** (cm^-1^) ν_max_: 1700 (CO); 3404 (NH). Yield = 18 %.

### Spectral data for 2-(2,4-dimethylphenoxy)-N-(4-iodo-2-methylphenyl)acetamide HA22

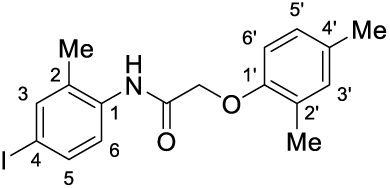

**^1^H-NMR** (400 MHz, CDCl_3_): δ 2.21 (3H, s, C^2^<u>Me</u>); 2.28 (3H, s, C^4’^<u>Me</u>); 2.31 (3H, s, C^2’^<u>Me</u>); 4.59 (2H, s, OCH_2_); 6.72 (1H, d, J=8.2 Hz, C^6’^H); 6.99 (1H, d, J=8.2 Hz, C^5’^H); 7.02 (1H, s, C^3’^H); 7.53 (1H, s, C^3^H); 7.54 (1H, d, J=8.7 Hz, C^6^H); 7.91 (1H, d, J=8.7 Hz, C^5^H); 8.31 (1H, br. s, NH). **^13^C-NMR** (100 MHz, CDCl_3_): δ 16.4 (C^2’^<u>Me</u>); 17.1 (C^2^<u>Me</u>); 20.5 (C^4’^<u>Me</u>); 67.7 (OCH_2_); 88.7 (C^4^I); 111.4 (C^6’^H); 123.3 (C^5^H); 125.9 (C^2’^); 127.5 (C^5’^H); 129.7 (C^2^); 131.6 (C^4’^); 132.1 (C^3’^H); 135.0 (C^1^); 136.0 (C^6^H); 139.0 (C^3^H); 152.9 (C^1’^O); 166.4 (NHCO). **MS** (ESI): m/z (%): 396 (M+H^+^, 100); 418 (M+Na^+^, 10). **IR** (cm^-1^) ν_max_: 1701 (CO); 2914; 3408 (NH). Yield = 14 %.

### Spectral data for 2-(4-chloro-2-methylphenoxy)-N-(4-fluoro-2-methylphenyl)acetamide HA23

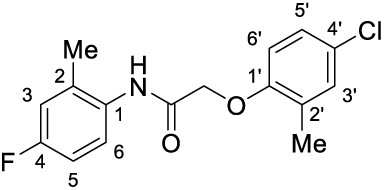

**^1^H-NMR** (400 MHz, CDCl_3_): δ 2.25 (3H, s, C^2^<u>Me</u>); 2.32 (3H, s, C^2’^<u>Me</u>); 4.61 (2H, s, OCH_2_); 6.76 (1H, d, J=8.6 Hz, C^6’^H); 6.88-6.97 (2H, m, C^3^H + C^5^H); 7.16 (1H, d, J=8.6 Hz, C^5’^H); 7.19 (1H, s, C^3’^H); 7.88-7.95 (1H, m, C^6^H); 8.14 (1H, br. s, NH). **^13^C-NMR** (100 MHz, CDCl_3_): δ 16.4 (C^2’^<u>Me</u>); 17.7 (C^2^<u>Me</u>); 67.8 (OCH_2_); 112.6 (C^6’^H); 113.5 (d, J^CF^=22.1 Hz, C^3^H); 117.1 (d, J^CF^=22.7 Hz, C^6^H); 124.1 (d, J^CF^=8.6 Hz, C^5^H); 127.0 (C^5’^H); 127.1 (C^2’^); 128.2 (C^4’^Cl); 130.7 (d, J^CF^=2.8 Hz, C^1^); 131.0 (d, J^CF^=7.8 Hz, C^2^); 131.1 (C^3’^H); 153.7 (C^1’^O); 159.9 (d, J^CF^=244.8 Hz, C^4^F); 165.9 (NHCO). **^19^F-NMR** (376 MHz, CDCl_3_): δ −116.9 (1F, m, C^4^F) **MS** (ESI): m/z (%): 308 (M+H^+^, 100); 330 (M+Na^+^, 30). **IR** (cm^-1^) ν_max_: 1668 (CO); 3404 (NH). Yield = 49 %.

### Spectral data for 2-(4-chloro-2-methylphenoxy)-N-(4-methoxy-2-methylphenyl)acetamide HA24

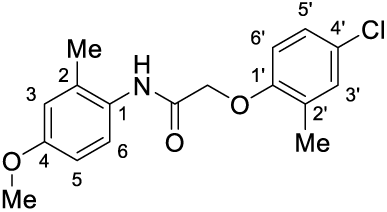

**^1^H-NMR** (400 MHz, CDCl_3_): δ 2.23 (3H, s, C^2^<u>Me</u>); 2.32 (3H, s, C^2’^<u>Me</u>); 3.78 (3H, s, O<u>Me</u>); 4.60 (2H, s, OCH_2_); 6.76 (1H, s, C^3^H); 6.76-6.82 (2H, m, C^6^H + C^6’^H); 7.16 (1H, d, J=8.6 Hz, C^5’^H); 7.18 (1H, s, C^3’^H); 7.77 (1H, d, J=8.4 Hz, C^5^H); 8.06 (1H, br. s, NH). **^13^C-NMR** (100 MHz, CDCl_3_): δ 16.4 (C^2’^<u>Me</u>); 17.9 (C^2^<u>Me</u>); 55.4 (OMe); 67.8 (OCH_2_); 111.7 (C^6^H); 112.6 (C^6’^H); 116.2 (C^3^H); 124.3 (C^5^H); 126.9 (C^1^); 127.0 (C^5’^H); 127.7 (C^2^); 128.2 (C^4’^Cl); 130.9 (C^3’^H); 131.0 (C^2’^); 153.8 (C^1’^O); 157.2 (C^4^O); 165.8 (NHCO). **MS** (ESI): m/z (%): 320 (M+H^+^, 100); 342 (M+Na^+^, 15). **HRMS** (ESI): Calculated for C_17_H_18_ClNO_3_^+^: 320.1047977 [M + H]^+^, found: 320.10369. **IR** (cm^-1^) ν_max_: 1683 (CO); 3415 (NH). Yield = 70 %.

### Spectral data for Methyl 4-(2-(4-chloro-2-methylphenoxy)acetamido)-3-methylbenzoate HA25

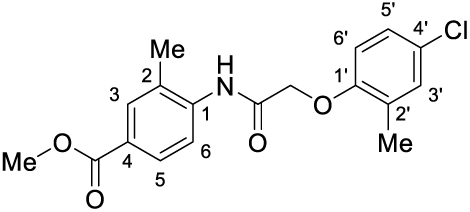

**^1^H-NMR** (400 MHz, CDCl_3_): δ 2.32 (3H, s, C^2^<u>Me</u>); 2.34 (3H, s, C^2’^<u>Me</u>); 3.90 (3H, s, O<u>Me</u>); 4.62 (2H, s, OCH_2_); 6.77 (1H, d, J=8.6 Hz, C^6’^H); 7.17 (1H, d, J=8.6 Hz, C^5’^H); 7.20 (1H, s, C^3’^H); 7.89 (1H, s, C^3^H); 7.92 (1H, d, J=8.5 Hz, C^5^H); 8.34 (1H, d, J=8.5 Hz, C^6^H); 8.44 (1H, br. s, NH). **^13^C-NMR** (100 MHz, CDCl_3_): δ 16.4 (C^2’^<u>Me</u>); 17.4 (C^2^<u>Me</u>); 52.1 (O<u>Me</u>); 67.7 (OCH_2_); 112.5 (C^6’^H); 120.2 (C^6^H); 126.1 (C^4^); 126.3 (C^1^); 127.0 (C^5’^H); 127.1 (C^4’^Cl); 128.1 (C^2’^); 128.9 (C^5^H); 131.1 (C^3’^H); 131.8 (C^3^H); 139.2 (C^2^); 153.5 (C^1’^O); 165.8 (NHCO). 166.6 (C^4^<u>C</u>O). **MS** (ESI): m/z (%): 348 (M+H^+^, 100); 370 (M+Na^+^, 30); 411 (20). **IR** (cm^-1^) ν_max_: 1666 (CO); 1697 (CO); 3406 (NH). Yield = 50 %.

### Spectral data for 2-(4-chloro-2-fluorophenoxy)-N-(4-iodo-2-methylphenyl)acetamide HA26

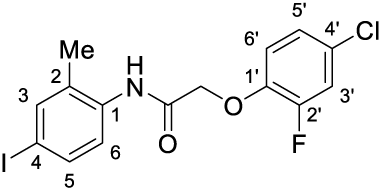

**^1^H-NMR** (400 MHz, CDCl_3_): δ 2.25 (3H, s, C^2^<u>Me</u>); 4.63 (2H, s, OCH_2_); 6.90-6.97 (1H, m, C^6’^H); 6.08-6.15 (1H, m, C^5’^H); 6.15-6.21 (1H, m, C^3’^H); 7.54 (1H, d, J=8.6 Hz, C^5^H); 7.55 (1H, s, C^3^H); 7.84 (1H, d, J=8.6 Hz, C^6^H); 8.37 (1H, br. s, NH). **^13^C-NMR** (100 MHz, CDCl_3_): δ 16.9 (C^2^<u>Me</u>); 68.8 (OCH_2_); 89.1 (C^4^I); 116.1 (d, J^CF^=2.0 Hz, C^6’^H); 117.5 (d, J^CF^=21.2 Hz, C^3’^H); 123.3 (C^6^H); 124.8 (d, J^CF^=3.8 Hz, C^5’^H); 127.8 (d, J^CF^=8.9 Hz, C^4’^Cl); 130.3 (C^2^); 134.7 (C^1^); 135.9 (C^5^H); 139.1 (C^3^H); 144.0 (d, J^CF^=11.0 Hz, C^1’^O); 152.3 (d, J^CF^=250.1 Hz, C^2’^F); 165.0 (NHCO). **^19^F-NMR** (376 MHz, CDCl_3_): δ −131.3 (1F, m, C^2’^F). **MS** (ESI): m/z (%): 420 (M+H^+^, 100); 442 (M+Na^+^, 70). **HRMS** (ESI): Calculated for C_15_H_12_ClFINO_2_^+^: 419.9658041 [M + H]^+^, found: 419.96545. **IR** (cm^-1^) ν_max_: 1699 (CO); 3404 (NH). Yield = 93 %.

### Spectral data for 2-(4-chloro-2-nitrophenoxy)-N-(4-iodo-2-methylphenyl)acetamide HA27

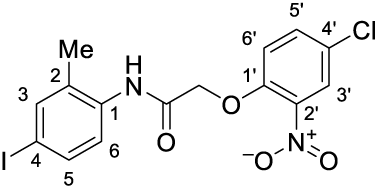

**^1^H-NMR** (400 MHz, CDCl_3_): δ 2.33 (3H, s, C^2^<u>Me</u>); 4.74 (2H, s, OCH_2_); 7.07 (1H, d, J=8.9 Hz, C^6’^H); 7.54 (1H, d, J=8.5 Hz, C^6^H); 7.57 (1H, s, C^3^H); 7.62 (1H, dd, J=8.9, 2.2 Hz, C^5’^H); 7.78 (1H, d, J=8.5 Hz, C^5^H); 8.07 (1H, d, J=2.2 Hz, C^3’^H); 8.56 (1H, br. s, NH). **^13^C-NMR** (100 MHz, CDCl_3_): δ 17.3 (C^2^<u>Me</u>); 68.0 (OCH_2_); 89.6 (C^4^I); 115.8 (C^6’^H); 124.1 (C^5^H); 126.7 (C^3’^H); 127.4 (C^4’^Cl); 131.4 (C^2^); 134.6 (C^1^); 135.3 (C^5’^H); 135.7 (C^6^H); 138.9 (C^2’^); 139.3 (C^3^H); 149.3 (C^1’^O); 164.1 (NHCO). **MS** (ESI): m/z (%): 447 (M+H^+^, 30); 469 (M+Na^+^, 100). **HRMS** (ESI): Calculated for C_15_H_12_ClIN_2_O_4_^+^: 446.9603041 [M + H]^+^, found: 446.96169. **IR** (cm^-1^) ν_max_: 1697 (CO); 3388 (NH). Yield = 52 %.

### Spectral data for 2-(4-chloro-2-cyclohexylphenoxy)-N-(4-iodo-2-methylphenyl)acetamide HA28

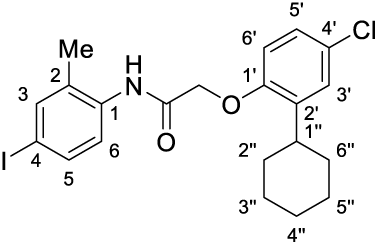

**^1^H-NMR** (400 MHz, CDCl_3_): δ 1.19-1.97 (10H, m, C^2’^CH(C<u>H</u>_2_)_5_); 2.20 (3H, s, C^2^<u>Me</u>); 2.90-3.00 (1H, tt, J=17.2, 2.9 Hz, C^2’^C^1’’^<u>H</u>); 4.61 (2H, s, OCH_2_); 6.78 (1H, d, J=8.7 Hz, C^6’^H); 7.13 (1H, dd, J=8.7, 2.6 Hz, C^5’^H); 7.23 (1H, d, J=2.6 Hz, C^3’^H); 7.53 (1H, s, C^3^H); 7.54 (1H, d, J=9.2 Hz, C^5^H); 7.81 (1H, d, J=9.2 Hz, C^6^H); 8.09 (1H, br. s, NH). **^13^C-NMR** (100 MHz, CDCl_3_): δ 17.3 (C^2^<u>Me</u>); 26.1 (C^4’’^H_2_); 27.0 (C^3’’^H_2_ + C^5’’^H_2_); 33.2 (C^2’’^H_2_ + C^6’’^H_2_); 37.3 (C^1’’^H); 68.3 (OCH_2_); 89.2 (C^4^I); 113.3 (C^6’^H); 123.9 (C^6^H); 126.7 (C^5’^H); 127.6 (C^3’^H); 127.8 (C^4’^Cl); 130.2 (C^2^); 134.7 (C^1^); 136.0 (C^5^H); 138.1 (C^2’^); 139.1 (C^3^H); 152.6 (C^1’^O); 166.0 (NHCO). **MS** (ESI): m/z (%): 225 (85); 282 (85); 484 (M+H^+^, 100). **HRMS** (ESI): Calculated for C_21_H_23_ClINO_2_^+^: 484.0534762 [M + H]^+^, found: 484.05214. **IR** (cm^-1^) ν_max_: 1703 (CO); 2922; 3412 (NH). Yield = 44 %.

### Spectral data for 2-(4-chloro-2-methoxyphenoxy)-N-(4-iodo-2-methylphenyl)acetamide HA29

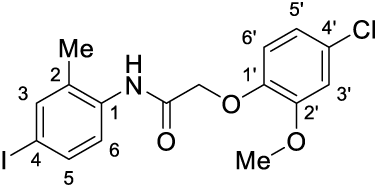

**^1^H-NMR** (400 MHz, CDCl_3_): δ 2.26 (3H, s, C^2^<u>Me</u>); 3.87 (3H, s, O<u>Me</u>); 4.60 (2H, s, OCH_2_); 6.84 (1H, m, C^5’^H); 6.89-6.93 (2H, m, C^3’^H + C^6’^H); 7.53 (1H, d, J=9.2 Hz, C^6^H); 7.54 (1H, s, C^3^H); 7.86 (1H, d, J=9.2 Hz, C^5^H); 8.63 (1H, br. s, NH). **^13^C-NMR** (100 MHz, CDCl_3_): δ 16.9 (C^2^<u>Me</u>); 56.0 (O<u>Me</u>); 69.3 (OCH_2_); 88.8 (C^4^I); 112.7 (C^3’^H); 115.6 (C^5’^H); 120.7 (C^6’^H); 123.4 (C^5^H); 128.1 (C^4’^Cl); 130.2 (C^2^); 135.0 (C^1^); 135.9 (C^6^H); 139.1 (C^3^H); 145.5 (C^2’^); 150.1 (C^1’^O); 166.0 (NHCO). **MS** (ESI): m/z (%): 432 (M+H^+^, 100); 454 (M+Na^+^, 40). **HRMS** (ESI): Calculated for C_16_H_15_ClINO_3_^+^: 431.9857906 [M + H]^+^, found: 431.98692. **IR** (cm^-1^) ν_max_: 1701 (CO); 3387 (NH). Yield = 74 %.

### Spectral data for 2-(2-benzyl-4-chlorophenoxy)-N-(4-iodo-2-methylphenyl)acetamide HA30

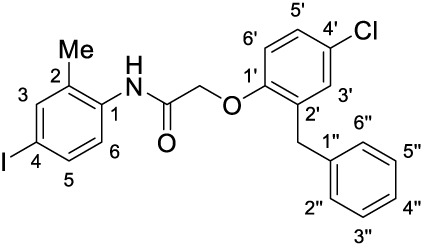

**^1^H-NMR** (400 MHz, CDCl_3_): δ 1.86 (3H, s, C^2^<u>Me</u>); 4.03 (2H, s, C^2’^CH_2_); 4.58 (2H, s, OCH_2_); 6.81 (1H, d, J=8.7 Hz, C^6’^H); 7.11-7.17 (3H, m, C^5’^H + C^3’’^H + C^5’’^H); 7.18 (1H, d, J=2.6 Hz, C^3’^H); 7.20-7.26 (3H, m, C^2’’^H + C^4’’^H + C^6’’^H); 7.44-7.53 (3H, m, C^3^H + C^5^H + C^6^H); 7.59 (1H, br. s, NH). **^13^C-NMR** (100 MHz, CDCl_3_): δ 16.9 (C^2^<u>Me</u>); 36.5 (C^2’^<u>C</u>H_2_); 67.8 (OCH_2_); 89.9 (C^4^I); 113.0 (C^6’^H); 124.9 (C^6^H); 126.6 (C^3’’^H + C^5’’^H); 127.4 (C^4’^Cl); 127.8 (C^2’’^H + C^6’’^H); 128.8 (C^4’’^H); 131.0 (C^2’^); 131.2 (C^3’^H); 131.9 (C^2^); 134.3 (C^1^); 135.8 (C^5^H); 139.1 (C^3^H); 139.3 (C^1’’^H); 153.5 (C^1’^O); 165.9 (NHCO). **MS** (ESI): m/z (%): 492 (M+H^+^, 100); 514 (M+Na^+^, 75). **HRMS** (ESI): Calculated for C_22_H_19_ClINO_2_^+^: 492.0221761 [M + H]^+^, found: 492.02355. **IR** (cm^-1^) ν_max_: 1670 (CO); 3253 (NH). Yield = 58 %.

### Spectral data for 2-(4-chloro-2-(isoxazole-5-yl)phenoxy)-N-(4-iodo-2-methylphenyl)acetamide HA31

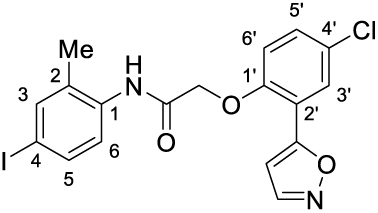

**^1^H-NMR** (400 MHz, CDCl_3_): δ 2.10 (3H, s, C^2^<u>Me</u>); 4.77 (2H, s, OCH_2_); 6.72 (1H, d, J=1.5 Hz, NC<u>H</u>); 6.99 (1H, d, J=8.9 Hz, C^6’^H); 7.43 (1H, dd, J=8.9, 2.5 Hz, C^5’^H); 7.52 (1H, s, C^3^H); 7.53 (1H, d, J=9.0 Hz, C^5^H); 7.66 (1H, d, J=9.0 Hz, C^6^H); 7.82 (1H, d, J=2.5 Hz, C^3’^H); 8.18 (1H, br. s, NH); 8.34 (1H, d, J=1.5 Hz, NCHC<u>H</u>). **^13^C-NMR** (100 MHz, CDCl_3_): δ 17.1 (C^2^<u>Me</u>); 68.2 (OCH_2_); 89.9 (C^4^I); 102.9 (N<u>C</u>H); 114.2 (C^6’^H); 118.2 (C^2’^); 124.6 (C^6^H); 128.0 (C^4’^Cl); 129.0 (C^3’^H); 131.4 (C^2^); 131.5 (C^5’^H); 134.4 (C^1^); 135.8 (C^5^H); 139.3 (C^3^H); 150.7 (NCH<u>C</u>H); 152.2 (C^1’^O); 165.0 (C^2’^<u>C</u>); 165.2 (NHCO). **MS** (ESI): m/z (%): 469 (M+H^+^, 100); 491 (M+Na^+^, 95). **HRMS** (ESI): Calculated for C_18_H_14_ClIN_2_O_3_^+^: 468.9810395 [M + H]^+^, found: 468.98041. **IR** (cm^-1^) ν_max_: 1668 (CO); 3244 (NH). Yield = 57 %.

### Spectral data for 2-(4-chloro-2-(1H-pyrazool-5-yl)phenoxy)-N-(4-iodo-2-methylphenyl)acetamide HA32

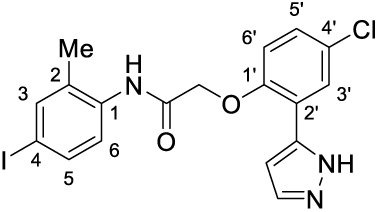

**^1^H-NMR** (400 MHz, CDCl_3_): δ 2.07 (3H, s, C^2^<u>Me</u>); 4.73 (2H, s, OCH_2_); 6.69 (1H, br. s, NCHC<u>H</u>); 6.93 (1H, d, J=8.8 Hz, C^6’^H); 7.31 (1H, dd, J=8.8, 1.9 Hz, C^5’^H); 7.40 (1H, d, J=8.2 Hz, C^6^H); 7.51 (1H, d, J=8.2 Hz, C^5^H); 7.53 (1H, s, C^3^H); 7.62 (1H, br. s, NC<u>H</u>); 7.66 (1H, d, J=1.9 Hz, C^3’^H); 9.37 (1H, br. s, NH). **^13^C-NMR** (100 MHz, CDCl_3_): δ 17.3 (C^2^<u>Me</u>); 68.4 (OCH_2_); 90.4 (C^4^I); 105.3 (NCH<u>C</u>H); 114.5 (C^6’^H); 126.1 (C^6^H); 127.6 (C^4’^Cl); 129.1 (C^5’^H); 129.4 (C^3’^H); 131.6 (N<u>C</u>H); 133.4 (C^2^); 134.8 (C^1^); 135.7 (C^5^H); 139.4 (C^3^H); 146.2 (C^2’^<u>C</u>); 152.7 (C^1’^O); 166.5 (NHCO). **MS** (ESI): m/z (%): 468 (M+H^+^, 100); 490 (M+Na^+^, 70). **HRMS** (ESI): Calculated for C_18_H_15_ClIN_3_O_2_^+^: 467.997024 [M + H]^+^, found: 467.99628. **IR** (cm^-1^) ν_max_: 1672 (CO); 2910 (NH); 3155 (NH). Yield = 74 %.

### Spectral data for Methyl 2-(2-(4-chloro-2-methylphenoxy)acetamido)-5-iodobenzoate HA33

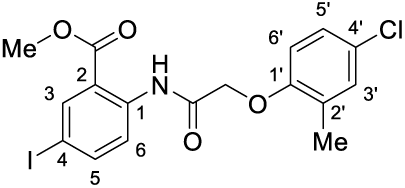

**^1^H-NMR** (400 MHz, CDCl_3_): δ 2.43 (3H, s, C^2’^<u>Me</u>); 3.89 (3H, s, O<u>Me</u>); 4.61 (2H, s, OCH_2_); 6.73 (1H, d, J=8.7 Hz, C^6’^H); 7.11 (1H, dd, J=8.7, 2.5 Hz, C^5’^H); 7.18 (1H, d, J=2.5 Hz, C^3’^H); 7.83 (1H, dd, J=8.9, 2.1 Hz, C^5^H); 8.33 (1H, d, J=2.1 Hz, C^3^H); 8.58 (1H, d, J=8.9 Hz, C^6^H); 11.81 (1H, br. s, NH). **^13^C-NMR** (100 MHz, CDCl_3_): δ 16.3 (C^2’^<u>Me</u>); 52.5 (O<u>Me</u>); 68.1 (OCH_2_); 85.9 (C^4^I); 112.2 (C^6’^H); 117.7 (C^2^); 122.6 (C^6^H); 126.5 (C^5’^H); 126.6 (C^4’^Cl); 129.4 (C^2’^); 131.0 (C^3’^H); 139.3 (C^3^H); 139.9 (C^1^); 154.0 (C^1’^O); 166.7 (C^2^<u>C</u>O); 167.3 (NHCO). **MS** (ESI): m/z (%): 225 (50); 256 (30); 371 (90); 447 (35); 460 (M+H^+^, 100); 482 (M+Na^+^, 30). **HRMS** (ESI): Calculated for C_17_H_15_ClINO_4_^+^: 459.9807052 [M + H]^+^, found: 459.97997. **IR** (cm^-1^) ν_max_: 1687 (CO); 3255 (NH). Yield = 90 %.

### Spectral data for 2-(4-chloro-2-methylphenoxy)-N-(4-iodo-2-nitrophenyl)acetamide HA34

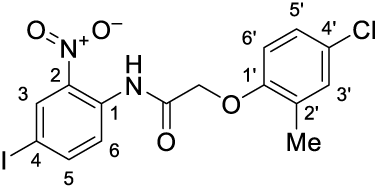

**^1^H-NMR** (400 MHz, CDCl_3_): δ 2.42 (3H, s, C^2’^<u>Me</u>); 4.65 (2H, s, OCH_2_); 6.74 (1H, d, J=8.7 Hz, C^6’^H); 7.14 (1H, dd, J=8.7, 2.5 Hz, C^5’^H); 7.21 (1H, d, J=2.5 Hz, C^3’^H); 7.96 (1H, dd, J=8.9, 2.1 Hz, C^5^H); 8.56 (1H, d, J=2.1 Hz, C^3^H); 8.68 (1H, d, J=8.9 Hz, C^6^H); 11.29 (1H, br. s, NH). **^13^C-NMR** (100 MHz, CDCl_3_): δ 16.4 (C^2’^<u>Me</u>); 67.7 (OCH_2_); 85.7 (C^4^I); 112.1 (C^6’^H); 123.8 (C^6^H); 126.6 (C^5’^H); 127.1 (C^4’^Cl); 129.1 (C^2’^); 131.3 (C^3’^H); 133.4 (C^2^); 134.2 (C^3^H); 137.0 (C^1^); 144.5 (C^5^H); 153.5 (C^1’^O); 167.3 (NHCO). **MS** (ESI): m/z (%): 225 (45); 280 (100); 302 (45); 371 (75); 393 (40); 447 (M+H^+^, 15). **IR** (cm^-1^) ν_max_: 1699 (CO); 3317; 3410 (NH). Yield = 43 %.

## Notes

### Competing Interest Statement

The authors have declared no competing interest.

